# Concurrent temporal patterning of neural stem cells in the fly visual system

**DOI:** 10.1101/2022.10.13.512100

**Authors:** Urfa Arain, Ishrat Maliha Islam, Priscilla Valentino, Ted Erclik

**Author notes:** These authors contributed equally (Urfa Arain, Ishrat Maliha Islam, Priscilla Valentino).

## Abstract

The temporal and spatial patterning of neural stem cells is a powerful mechanism by which to generate neural diversity in both vertebrate and invertebrate brains. In the *Drosophila* optic lobe, the neuroblasts (NBs) that generate the ∼120 neuronal cell types of the medulla are patterned by independent temporal and spatial inputs. In the temporal axis, a cascade of twelve transcription factors (TFs) are expressed in medulla NBs as they age. In the spatial axis, the neuroepithelium from which these NBs are generated is sub-divided into eight compartments by the expression of five additional TFs. Distinct neuronal types are generated by NBs based on their spatio-temporal address. Here, we describe a third major patterning axis that further diversifies neuronal fates in the medulla. We show that the symmetrically dividing neuroepithelial cells from which the medulla NBs are generated are temporally patterned by opposing gradients of the Imp and Syp RNA-binding proteins. Imp and Syp regulate the expression of a set of TFs in the neuroepithelium to confer NBs from the same spatio-temporal address with unique identities based on the developmental stage they are generated. We show that Imp and Syp differentially pattern NBs in the Vsx1-Hth spatio-temporal birth window to generate seven distinct neuronal cell types (Li2, TmY17, TmY15, Tm23, Pm3a, Pm3b and TmY12) in successive developmental windows. We further demonstrate that the birthdate of these neurons correlates with their final position in the adult cortex, resulting in unanticipated specializations of the retinotopic circuit in the anterior-posterior axis of the visual system. The concurrent temporal patterning of symmetrically and asymmetrically dividing neural stem cells thus acts as a powerful mechanism to couple the generation of neural diversity with circuit patterning.

## Introduction

The development of complex neural circuits requires that stem cells generate diverse types of neurons in the correct number and location. Pioneering work in insects and vertebrates has shown that the spatial and temporal patterning of neural stem cells plays a critical role in regulating multiple aspects of neurogenesis. In the *Drosophila* embryonic ventral nerve cord, neural stem cells, termed neuroblasts (NBs), integrate spatial and temporal inputs at both the transcriptional and epigenetic levels to generate neural diversity (Isshiki et al. 2001; Karlsson et al. 2010; Sen et al. 2019; Rossi et al. 2021). In the spatial axis, patterning by segment polarity, columnar and Hox genes assign each NB a unique positional identity along the dorsal-ventral (D-V) and anterior-posterior (A-P) axes of the embryo (Skeath et al. 1995; Hirth et al. 1998; McDonald et al. 1998; Bhat 1999; Lin and Lee 2012). In the temporal axis, NBs sequentially express a series of temporal transcription factors (tTFs) - Hunchback, Kruppel, Pdm1/2, Castor, and Grainyhead - as they age to generate distinct neuronal types in a birth order-dependent manner (Kambadur et al. 1998; Isshiki et al. 2001; Doe 2017). Temporal and spatial stem cell patterning has been observed in multiple vertebrate brain regions as well, including the neural tube, where localized morphogen activity assigns unique positional identities to neural stem cells and distinct sets of tTFs are expressed in neurons based on their developmental birth order (Marti et al. 1995; Briscoe et al. 2000; Lee et al. 2000; Dasen et al. 2003; Delile et al. 2019; Sagner and Briscoe 2019; Sagner et al. 2021). The temporal patterning of vertebrate neural stem cells has been particularly well studied in the developing neuroretina, where a suite of tTFs, including Ikaros and Casz1, homologs of the *Drosophila* tTFs Hunchback and Castor, regulate the competence of progenitor cells to generate retinal cell-types in a birth order-dependent manner (Elliott et al. 2008; Mattar et al. 2015; Clark et al. 2019; Javed et al. 2020).

In recent years, the *Drosophila* medulla has emerged as a powerful model system in which to study how neural stem cells integrate spatial and temporal patterning inputs to develop a complex neural circuit. The medulla, which together with the lamina and lobula complex comprises the optic lobe, mediates the processing of both color and motion information (Fischbach and Dittrich 1989; Hofbauer and Campos-Ortega 1990; Morante and Desplan 2008; Behnia et al. 2014; Nériec and Desplan 2016; Kind et al. 2021). Its 40,000 cells are organized into 800 repeating columns that propagate the retinotopic inputs received from the overlying ommatidia (Nériec and Desplan 2016; Millard and Pecot 2018). The ∼120 neuronal cell types of the medulla develop from a neuroepithelial (NE) crescent termed the outer proliferation center (OPC) (Egger et al. 2007; Yasugi et al. 2008; Li et al. 2013; Apitz and Salecker 2014; Erclik et al. 2017; Özel et al. 2020). Beginning at the onset of the third larval instar, and continuing for three days, a proneural wave expands in a medial-to-lateral direction to convert NE cells into NBs (Ceron et al. 2001; Yasugi et al. 2008; Egger et al. 2010; Yasugi et al. 2010). In the wake of the proneural wave, the youngest NBs are located laterally, closest to the OPC NE, whereas the oldest NBs are found medially, adjacent to the central brain (Yasugi et al. 2008; Egger et al. 2010; Yasugi et al. 2010). Once specified, NBs divide asymmetrically multiple times to generate intermediate Ganglion Mother Cells (GMCs), which then divide once more to produce the neurons and glia of the medulla (Ceron et al. 2001; Egger et al. 2007).

The integration of spatial and temporal inputs contributes to the generation of neuronal diversity in the medulla (Bertet et al. 2014; Erclik et al. 2017). In the spatial axis, the OPC NE from which the NBs are derived is patterned by the expression of three homeobox TFs: Vsx1 in the center of the crescent, Optix in the arms, and Rx at the tips (Erclik et al. 2017) (Fig. 1A). The Rx region is further sub-divided by the expression of the signaling molecules Dpp and Wingless, and the dorsal and ventral halves of the OPC are defined by two zinc finger TFs, Spalt and Disco (Kaphingst and Kunes 1994; Valentino and Erclik 2022). In the temporal axis, a cascade of twelve tTFs - including the sequentially expressed TFs Homothorax (Hth), Eyeless (Ey), Sloppy-paired 1/2 (Slp1/2), Dichaete (D), and Tailless (Tll) - pattern NBs as they age (Hasegawa et al. 2011; Li et al. 2013; Konstantinides et al. 2022; Zhu et al. 2022) (Fig. 1A). Distinct neuronal cell-types are generated by NBs based on their spatio-temporal address (Erclik et al. 2017). For example, Pm3 neurons are generated by NBs derived from the Vsx1 spatial compartment and Hth temporal window (Erclik et al. 2017).

**Figure 1:**
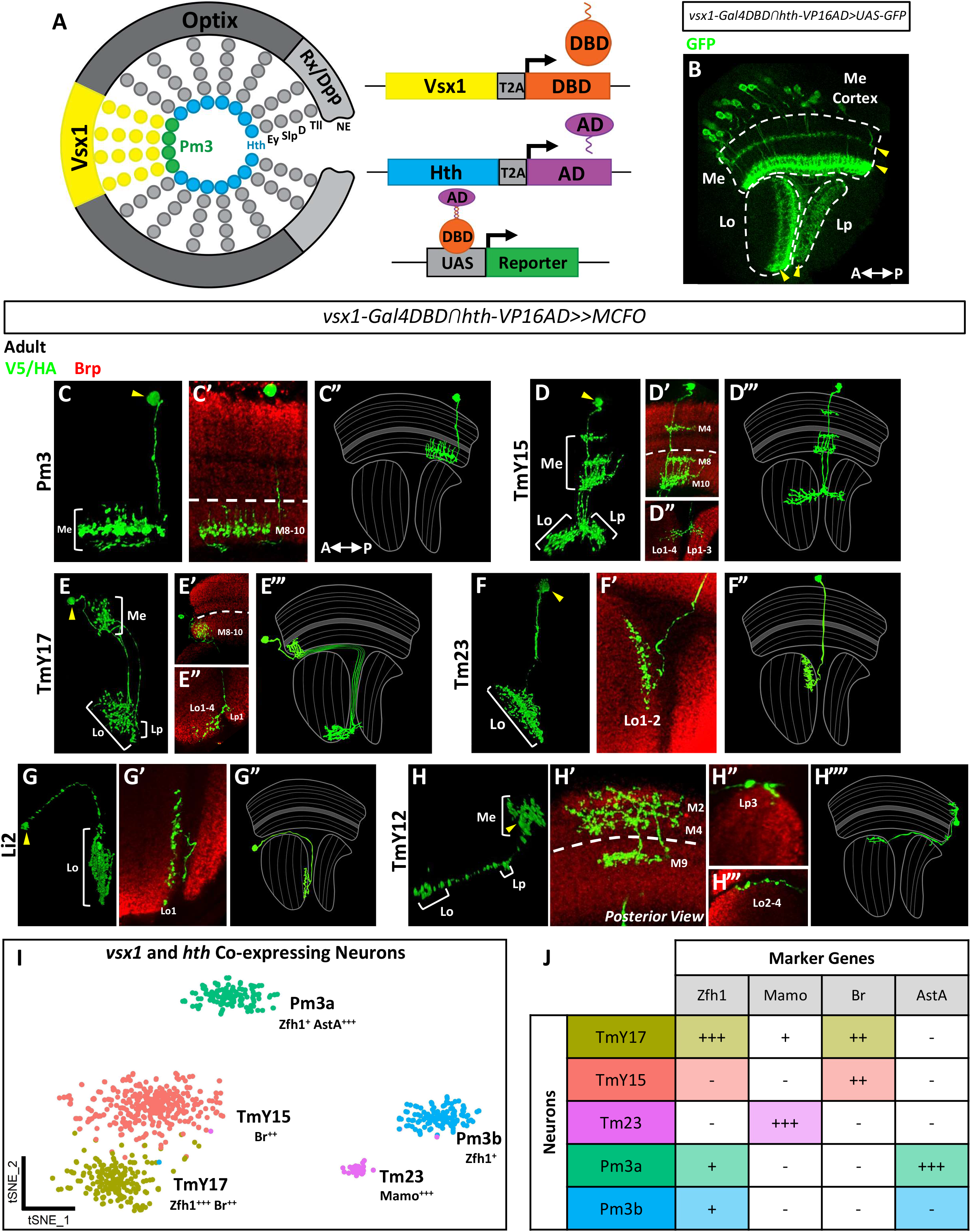
Seven distinct neuronal types are labeled by the vsx1ꓵhth split-Gal4 driver. **A:** Schematic of the split-Gal4 system used to drive expression of a reporter in Vsx1 and Hth co-expressing neurons. Neurons born in the Vsx1 spatial compartment (yellow) express the Gal4 DNA-binding domain (DBD) and neurons born from the Hth temporal window (blue) express the Gal4 transcription activation domain (AD). Gal4 transcriptional activity is reconstituted in Vsx1 and Hth co-expressing neurons (green), leading to the expression of a UAS-driven reporter. **B:** GFP (green) expression in the adult optic lobe driven by the *vsx1ꓵhth-Gal4* driver. The medulla (Me), lobula (Lo), and lobula plate (Lp) are indicated with white dashed lines. Yellow arrowheads denote labeled neuropil layers with unexpected innervations, not contributed to by Pm3 neurons. Anterior is left. **C-H:** MCFO clones (V5/HA, green) and schematics of the 6 neuron types labeled by *vsx1ꓵhth-Gal4* in the adult optic lobe neuropils (Brp, red). (**C**) Pm3, (**D**) TmY15, (**E**) TmY17, (**F**) Tm23, (**G**) Li2, and (**H**) TmY12. 3D reconstructions (**C, D, E, F, G, H**), single confocal sections of medulla arborizations (**C’, D’, E’, H’**), single confocal sections of lobula/lobula plate arborizations (**D”, E”, F’, G’, H”, H’’’**) and 2D whole-neuron schematics (**C’’, D’’’, E’”, F”, G”, H’’’’**) are shown for each neuron. White dashed lines indicate the M7 medulla layer and yellow arrowheads indicate the cell body. In all images (except **H’**, as indicated) anterior is left. **I**: tSNE visualization of *vsx1* and *hth* co-expressing cells subsetted from a whole adult *Drosophila* optic lobe single-cell RNAseq dataset. Neuronal clusters are annotated with cell type identities in accordance with marker analysis performed using *vsx1ꓵhth-Gal4*. Marker combinations used to annotate the neuronal clusters are also indicated. **J**: Table summarizing the expression profiles of marker genes, as identified through immunohistochemistry analysis, in neurons labeled by *vsx1ꓵhth-Gal4*.

Here we describe a third patterning axis that operates together with the previously reported spatial and temporal axes to generate neuronal diversity in the medulla. We demonstrate that opposing gradients of the RNA-binding proteins, IGF-II mRNA-binding protein (Imp) and Syncrip (Syp), temporally pattern the NE from which the NBs are derived. Imp and Syp regulate the downstream TFs Chronologically inappropriate morphogenesis (Chinmo), Maternal gene required for meiosis (Mamo), and Ecdysone-induced protein 93F (E93) to assign NBs unique temporal identities based on when during development they are generated by the proneural wave. As proof of principle, we show that Imp and Syp patterning diversifies the progeny of Vsx1-Hth NBs; in addition to the previously reported Pm3 neurons, six additional cell types are generated at this spatial-temporal address. We further show that these neurons occupy stereotyped positions along the A-P axis of the adult medulla in accordance with their developmental birth order; earlier-born neurons are located anteriorly, whereas later-born neurons are located posteriorly. We find that two of these cell types, TmY17 and TmY12, exclusively innervate the medulla columns located closest to their cell bodies, which results in unanticipated patterning of the fly’s retinotopic circuit along the A-P axis. The concurrent temporal patterning of symmetrically and asymmetrically dividing neural stem cells thus acts as a powerful mechanism to couple neurogenesis with circuit assembly.

## Results

### Unexpected neuronal diversity at the intersection of the Vsx1 spatial and Hth temporal axes

Previous work has shown that Pm3 neurons are born at the intersection of the Vsx1 spatial and Hth temporal axes and uniquely co-express these two TFs throughout their development (Erclik et al. 2017). To allow for the visualization and genetic manipulation of developing Pm3 neurons, we generated a split-Gal4 line (Luan et al. 2006) comprised of the *vsx1-Gal4DBD* and *hth-VP16AD* transgenes (Figure 1A). We confirmed that expression of the *vsx1⋂hth-Gal4* is restricted to cells that co-express Vsx1 and Hth by crossing the driver to *UAS-GFP* and co-staining larval and adult brains with Vsx1 and Hth antibodies (Fig. S1A-B). In the 3^rd^ instar larva, *vsx1⋂hth-Gal4>GFP* expression is restricted to the NE cells, NBs, and neurons of the cOPC in which Vsx1 and Hth are co-expressed (Fig. S1A). In the adult medulla cortex, we observed a perfect overlap between the neurons that co-express Vsx1, Hth, and GFP, indicating that the split-Gal4 is a faithful reporter for cells that co-express these two patterning factors (Fig. S1B). Surprisingly, however, we observed that the neuropil innervations of the neurons labeled by *vsx1⋂hth-Gal4>GFP* are more extensive than what would be expected from Pm3 neurons alone; in addition to the characteristic Pm3 arborizations located in the proximal medulla (Hasegawa et al. 2011; Erclik et al. 2017), *vsx1⋂hth-Gal4>GFP* labeled neurons also innervate the distal medulla, lobula, and lobula plate neuropils (Fig. 1B). The unexpected complexity of the neuropil arborizations labeled by this driver line suggests that other neuronal types, in addition to Pm3 neurons, are generated at the Vsx1-Hth spatio-temporal birth address.

To identify the cell types labeled by the *vsx1⋂hth-Gal4* driver, we used the MultiColor FlpOut (MCFO) system (Nern et al. 2015) to generate single-cell clones of neurons in the adult brain. Remarkably, we found that 6 morphologically distinct neuronal cell types are labeled by *vsx1⋂hth-Gal4*: Pm3, TmY15, TmY17, Tm23, Li2, and TmY12 (Fig. 1C-H). Pm3 neurons, which represent the previously reported output of Vsx1-Hth NBs, are medulla-intrinsic neurons that arborize in the proximal M8-M10 layers of the medulla (Fig. 1C) (Hasegawa et al. 2011; Erclik et al. 2017). TmY15 and TmY17 neurons are similar in morphology in that they both arborize in the proximal medulla layers M8-M10 and send multiple axons to the lobula and lobula plate (Fig. 1D-E). However, TmY15 neurons additionally possess arborizations in distal medulla layer M4 and extend processes into deeper layers of the lobula plate (Lp1-3 for TmY15 vs Lp1 for Tmy17) (Fig. 1D-E). TmY15 neurons have been previously identified as components of the motion detection pathway (Takemura et al. 2017; Shinomiya et al. 2019), whereas TmY17 neurons have not been described. Tm23 neurons, which have also been characterized as components of the motion detection pathway, send their axons directly through the medulla and into the lobula, where they extend wide arborizations in Lo1-2 (Fig. 1F) (Fischbach and Dittrich 1989; Shinomiya et al. 2019). Li2 neurons are lobula intrinsic neurons that arborize in the superficial Lo1 layer (Fig. 1G) (Fischbach and Dittrich 1989). Finally, TmY12 neurons arborize in layers M2, M4, and M9 of the medulla and extend their axons into the lobula complex where they innervate both the lobula (Lo2-4) and lobula plate (Lp3) (Fig. 1H) (Fischbach and Dittrich 1989).

We next determined whether the 6 neuronal cell types labeled by *vsx1⋂hth-Gal4* are discoverable in a recently published single-cell RNAseq dataset of the adult optic lobe (Özel et al. 2020). We identified three clusters of cells (Pm3, 158, and 159) in the dataset which had significantly upregulated levels of *vsx1* and *hth* (Wilcoxon rank sum, P-adj<0.05). Subsetting and independent re-clustering of the cells in these identity classes led to the emergence of 5 distinct clusters: Pm3, 158a, 158b, 159a, and 159b (Fig. 1I and S1C). To assign neural identities to these clusters, we labeled cells of the *vsx1⋂hth-Gal4* driver line with cluster-specific marker combinations (Fig. 1J and S1D-R). Using this approach, we assigned the identity of Tm23 to cluster 158b, based on the predicted and observed expression of high levels of the TF, Mamo, in these cells (Fig. S1C-D and P). We assigned the identities of TmY15 and TmY17 to two closely related clusters, 159a and 159b, respectively (Fig. S1C). Both cell types express Broad (Br) (Fig. S1E and L-M), but cluster 159b/TmY17 can be distinguished by the additional expression of high levels of the zinc finger homeodomain 1 (Zfh1) TF and low levels of Mamo (Fig. S1D, H and L-O). Unexpectedly, we found that *vsx1⋂hth-Gal4* labels two types of Pm3 neurons: The first, marked by the expression of Allatostatin A (AstA) and E93, maps to the previously annotated Pm3 cluster (Özel et al. 2020) and is henceforth referred to as Pm3a (Fig. S1C, F, I, K, Q). The second, referred to as Pm3b, was assigned to cluster 158a via the expression of Seven-up (Svp) and absence of E93/AstA expression (Fig. S1C, F-G and I-J). These two Pm3 neuronal sub-types, which are morphologically very similar, can also be differentiated from the other *vsx1⋂hth-Gal4* neuron types by their shared expression of low levels of Zfh1 (Fig. S1R). The only neurons labeled by *vsx1⋂hth-Gal4* that we were unable to identify in the single-cell RNAseq dataset were Li2 and TmY12. We posit that these neurons are present in too low a number in the adult optic lobe to form identifiable clusters in the single-cell RNAseq dataset, as we infrequently generated clones of either cell type during our MCFO analysis. Taken together, the above clonal and single-cell RNAseq data demonstrate that not 1, but 7, neuronal types are generated at the intersection of the Vsx1 spatial and Hth temporal axes: Pm3a (AstA/E93/low Zfh1), Pm3b (Svp/low Zfh1), Tm23 (high Mamo), Tmy17 (high Zfh1/low Mamo/Br), TmY15 (Br), Li2 and TmY12. The unexpected neuronal diversity generated at this spatial-temporal address suggests that an unidentified mechanism exists (in addition to spatial and temporal patterning) to further diversify cell fates in the medulla.

### Vsx1-Hth neuronal cell types are sequentially born over the 3 days of medulla neurogenesis

In our clonal analysis of Vsx1-Hth neurons, we observed that cell bodies of the same neuronal type occupy similar positions along the A-P axis of the medulla cortex. To determine whether neurons of the same type are indeed asymmetrically distributed along the A-P axis, we used cell type-specific marker combinations to examine the cell body distributions of *vsx1⋂hth-Gal4* neurons in the medulla. Remarkably, we observed that while the cell bodies of *vsx1⋂hth-Gal4* cell types are distributed throughout the D-V axis, they occupy restricted regions along the A-P axis: TmY17 cells are located in the anterior region of the cortex, followed by (from anterior to posterior) the cell bodies of TmY15, Tm23, Pm3b and Pm3a neurons (Fig. 2A-B). While a lack of cell-type specific markers prevented us from mapping the cell body positions for Li2 and TmY12 neurons, two observations suggest that these neurons are located at the anterior and posterior margins of the cortex, respectively: (i) the cell bodies of MCFO-labeled Li2 and TmY12 neurons are consistently observed at the anterior and posterior edges of the medulla (n=5 for Li2 and n=7 for TmY12), and (ii) small clusters of 2-3 cells, which are negative for the above marker combinations, are located at the anterior and posterior margins of the cortex (Fig. 1G-H and 2A-B).

**Figure 2:**
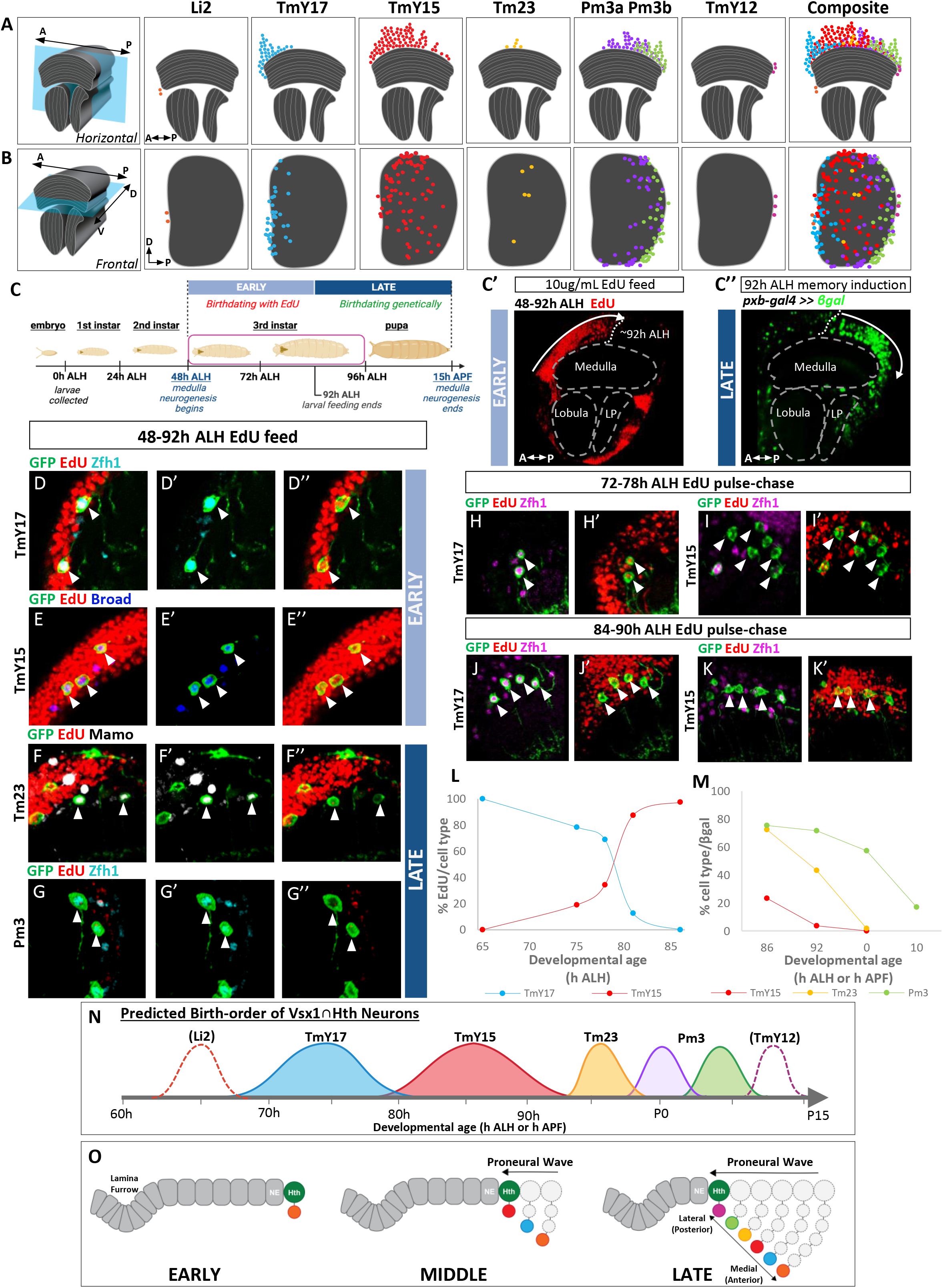
vsx1ꓵhth neuronal cell bodies occupy distinct A-P positions in the adult medulla cortex which correlate to a sequential birth-order in the larva. **A-B**: Schematic depicting cell body positions of *vsx1ꓵhth* neuronal cell-types in the medulla cortex in horizontal (**A**) and frontal (**B**) orientations. **A:** Horizontal views of *vsx1ꓵhth-Gal4* neurons in the A-P axis, where anterior is left. Cell bodies occupy discrete regions along the A-P axis in the following order: Li2 cell bodies (orange) are located most anterior, followed by TmY17 (blue), TmY15 (red), Tm23 (yellow), Pm3a/Pm3b (green/purple), and TmY12 (pink). **B:** Frontal views of *vsx1ꓵhth-Gal4* neurons in both A-P and D-V axes, where anterior is left and dorsal is up. The most anterior (Li2) and posterior (TmY12) cell bodies are also restricted to specific regions along the D-V axes. Cell bodies of TmY17 (blue), TmY15 (red) and Tm23 (yellow) are dispersed throughout the D-V axis. **C**: Schematic displaying the timing of medulla neurogenesis relative to fly development. Medulla neurogenesis begins in the late 2^nd^ instar (∼48 h ALH) and continues into pupation (∼15 h APF). Medulla neurons generated during the larval feeding window were birthdated by EdU and classified as “early-born” (light blue text box). Neurons generated after the feeding window were birthdated genetically via heat shock-induced lineage-trace experiments and classified as “late-born” (dark blue text box). (**C’**) Early-born EdU-labeled cells (red) occupy the anterior half of the adult medulla cortex (arrow, dashed line). (**C”**) Later-born βgal^+^ cells (green), labeled by a memory induction after 92 h ALH, occupy the posterior half of the adult medulla cortex (dashed line, arrow). **D-G:** EdU administered throughout the entire larval feeding window (48-92 h ALH) to identify early-born *vsx1ꓵhth-Gal4* neurons. (**D-D”**) Strongly Zfh1^+^ TmY17 neurons (cyan) are labeled by EdU and are therefore early-born (white arrowheads). (**E-E”**) Broad^+^ TmY15 neurons (blue) are labeled by EdU and are also early-born (white arrowheads). (**F-F”**) Strongly Mamo^+^ Tm23 neurons (white) are not labeled by EdU and are therefore late-born (post 92h ALH; white arrowheads). (**G-G”**) Weakly Zfh1+ Pm3 neurons (cyan) are not labeled by EdU and are therefore also late-born (post 92 h ALH; white arrowheads). **H-K:** EdU administered in 6 h pulses during the larval feeding window shows that TmY17 neurons are born before TmY15 neurons. (**H-I’**) EdU pulse in early third-instar (72-78 h ALH) labels Zfh1^+^ TmY17 neurons (**H-H’**), but not Zfh1^-^ TmY15 neurons (**I-I’**) in the adult. (**J-K’**) EdU pulse in the late third-instar (84-90 h ALH) does not label Zfh1^+^ TmY17 neurons (**J-J’**), but does label Zfh1^-^ TmY15 neurons (**K-K’**). **L:** Graph representing the proportion of total EdU-labeled TmY17 neurons (blue curve) vs. TmY15 neurons (red curve) after short pulses of EdU administered at different developmental stages (see methods). TmY17 neurons are labeled in earlier larval stages (68-80 h ALH), whereas TmY15 neurons are mostly labeled in later larval stages (80 h ALH onwards). Thus, TmY17 neurons are born before TmY15 neurons (n=10 brains per developmental stage, Chi-square test, *P<0.05). **M:** Graph representing the proportion of a given cell type labeled by βgal after heat shock at different developmental stages. TmY15 neurons (red curve) continue to be labeled at an 86 h ALH memory induction, but are no longer labeled after 92 h ALH. Tm23 neurons (yellow curve) are labeled by a memory induction at 86 h ALH and 92 h ALH, but are no longer labeled at the onset of pupation (0 h APF). Pm3 neurons are labeled in all memory induction experiments (green curve), with a steep decline after pupation (0-10 h APF). Thus, Tm23 neurons are born after TmY15 neurons, followed by Pm3 neurons (n=12 brains per developmental stage, Chi-square test, *P<0.05). **N:** Schematic summarizing the predicted birth order of *vsx1ꓵhth-Gal4* neurons across developmental time. Birth order of *vsx1ꓵhth-Gal4* neurons correlates to anteroposterior cell body locations (as shown in **A**). Li2 and TmY12 neurons could not be birthdated due to a lack of cell type-specific markers but are assigned to developmental origins based on their A-P cell body locations (Li2; dashed orange outline and TmY12; dashed pink outline). **O:** Model illustrating the relationship between neuronal birth order and proneural wave progression during medulla neurogenesis. A cross-section of the Vsx1 epithelium (NE, grey) shows the proneural wave sequentially converting OPC NE cells to NBs in the medial to lateral direction during development. Neurons generated from the OPC NE at an early developmental stage are medially-located (anterior in the adult), whereas neurons generated later in development are laterally-located (posterior in the adult).

We next asked how the striking asymmetric distribution of Vsx1-Hth neurons across the adult A-P axis is established during development. Notably, the A-P axis of the adult medulla cortex corresponds to the medial-lateral axis of the larval OPC. During medulla neurogenesis, a proneural wave converts OPC NE cells into NBs in a medial to lateral direction (Yasugi et al. 2008; Yasugi et al. 2010). A consequence of this moving wave is that neurons born earlier in development occupy a medial position in the larval cortex, whereas later-born neurons are located laterally. We thus asked whether the unexpected asymmetric distribution of Vsx1-Hth neurons along the A-P axis in the adult cortex could be due to the sequential generation of these cell types as a product of the proneural wave.

Medulla neurogenesis spans a period of 3 days, with the proneural wave commencing at 60 h after larval hatching (ALH) in the 3^rd^ instar larva and ending at 15 h after puparium formation (APF) in the early pupa (Egger et al. 2007; Yasugi et al. 2008). We developed EdU birth-dating and heat shock-induced lineage techniques to determine when neurons are born during this three-day period of neurogenesis (Fig. 2C). We found that the rearing of larvae on EdU-containing food from as early as 48 h ALH labeled neurons born only during the first half of neurogenesis. The failure to label later-born neurons is likely because larvae stop feeding at ∼92 h ALH in anticipation of molting. Consequently, in the adult brains of larvae fed from 48-92 h ALH, EdU-positive cells occupy only the anterior half of the medulla cortex (Fig. 2C’). In shorter feeding intervals, the timing of EdU administration correlates to the A-P position of labeled cells in the adult cortex; earlier feeds label more anterior cells, whereas later feeds progressively label more posterior cells (Fig. S2A-B). To birthdate neurons born after 92 h ALH, we utilized a lineage-based clonal approach that permanently labels neurons generated after heat shock. In this system, heat shock-induced clones are restricted to the Vsx1-region using the NE driver, *pxb-Gal4*, to drive FLP-recombinase expression (Suzuki et al. 2013). FLP-mediated recombination subsequently results in the permanent labeling of cells with βgal via the excision of a stop cassette. Thus, neurons born after heat shock are labeled by βgal, whereas neurons born earlier are unlabeled. We found that heat shock at 92 h ALH led to the labeling of neurons in the posterior half of the adult medulla cortex, in a complementary pattern to what we observed with 48-92 h ALH EdU feeds (Fig. 2C”). Heat shocks at later time points progressively labeled fewer anterior cells in the adult cortex (Fig. S2C). Thus, a combination of EdU and lineage-based techniques allows for the birth-dating of neurons over the course of medulla neurogenesis.

We next used these techniques to determine the birthdate of Vsx1-Hth cell types. We fed EdU to *vsx1⋂hth-Gal4>GFP* larvae from 48-92 h ALH and found that only 2 of the 7 cell-types incorporate EdU: TmY15 and TmY17 (Fig. 2D-E). In contrast, Tm23, Pm3a, and Pm3b neurons are either unlabeled or only partially labeled by EdU, indicating that these neuronal types are generated after the larval feeding window ends at 92 h ALH (Fig. 2F-G). We next determined whether TmY15 and TmY17 neurons are generated in the same or distinct windows by administering shorter 6 h EdU pulses at different points between 60-92 h ALH (Fig. 2H-K). We found that TmY17 neurons are born first, as they incorporate EdU during 6 h feeds that span the interval between 68 h to 82 h ALH (Fig. 2L). In contrast, TmY15 EdU incorporation was only observed when larvae were fed at later intervals between 78 h and 92 h ALH (Fig. 2L). Our EdU birth-dating analysis thus demonstrates that the neuronal cell types generated at the intersection of the Vsx1-Hth axes are sequentially born over the course of neurogenesis; TmY17 neurons are generated first, followed by TmY15 neurons and, subsequently, Tm23/Pm3 neurons, which are generated in a late window that spans the interval from the late 3^rd^ instar to the end of neurogenesis in the early pupa.

To determine the birth windows for the late-born Tm23 and Pm3 neurons, we generated heat shock-induced lineage clones at four time points (86 h ALH, 92 h ALH, 0 h APF and 10 h APF) and determined which cell types are labeled with βgal in the adult brain. Heat shock induction at 86 h ALH labeled a small number of TmY15 neurons and the majority of Tm23 and Pm3 neurons (Fig. S2D-G). This finding is consistent with our EdU data that TmY15 neurons continue to be generated at this time point. In contrast, heat shock-induction at 92 h ALH failed to label TmY15 neurons but did label a subset of Tm23 neurons and the majority of Pm3 neurons, indicating that the transition between the TmY15 and later windows occurs at ∼92 h ALH (Fig. S2H-J). A 0 h APF heat shock resulted in the loss of Tm23, but not Pm3, labeling, which indicates that Tm23 neurons are generated in the window between 92 h ALH and 0 h APF (Fig. S2K-L). Finally, a 10 h APF heat shock continued to label Pm3 neurons, though fewer than observed with a 0 h APF heat shock, indicating that the Pm3 window likely extends from 0 h APF until the end of neurogenesis (Fig. S2M).

Taken together, our EdU and clonal birth-dating experiments demonstrate that Vsx1-Hth neuronal types are sequentially born during medulla neurogenesis. TmY17 neurons (68-80 h ALH) are generated first, followed by TmY15 (80-92 h ALH), Tm23 (92 h ALH - 0 h APF), and Pm3 (0-15 h APF) neurons (Fig. 2L, M and N). The sequential birth order of these neurons is thus correlated with their cell body positions along the A-P axis of the adult medulla, falling in line with the progression of the proneural wave in the medial-to-lateral direction during neurogenesis (Egger et al. 2007; Yasugi et al. 2008) (Fig. 2A-B and O). While our analysis was unable to discriminate between the two Pm3 sub-types, the A-P distribution of these neurons suggests that Pm3b neurons are generated before Pm3a (Fig. 2A-B and N). Similarly, the A-P cell body positions of Li2 and TmY12 neurons suggest that Li2 neurons are born earliest, whereas TmY12 neurons are generated last (Fig. 2A-B and N). These findings suggest that Vsx1-Hth NBs make distinct neuronal types based on when during the 3 days of neurogenesis they undergo the NE-NB transition. We next asked whether the temporal patterning of the OPC NE from which the NBs are derived could account for this additional neuronal diversity.

### The OPC NE is temporally patterned by opposing gradients of the Imp and Syp RNA-binding proteins

A search for factors that are differentially expressed in the NE over the course of medulla neurogenesis identified the Imp and Syp RNA-binding proteins (Fig. 3A and B). We found that Imp and Syp are temporally expressed in opposing gradients in the OPC NE (Fig. 3C). Imp is expressed at high levels in the NE at 48 h ALH and declines in a gradient manner over time, becoming undetectable at 96 h ALH (Fig. 3D). Conversely, Syp expression is absent at 48 h ALH and increases over the course of neurogenesis, resulting in high levels of expression at 96 h ALH and through to the end of neurogenesis (Fig. 3E and I). Opposing gradients of Imp and Syp have been previously described in the NB lineages of the central brain, where the two proteins act to diversify neuronal fates via the post-transcriptional regulation of downstream TFs (Zhu et al. 2006; Liu et al. 2015; Ren et al. 2017; Syed et al. 2017; Liu et al. 2019; Islam and Erclik 2022). In the mushroom body lineage, Imp and Syp diversify fates via the post-transcriptional regulation of the Chinmo and Mamo TFs in their neuronal progeny (Liu et al. 2015; Liu et al. 2019). We thus asked whether Chinmo and Mamo are also temporally expressed in the OPC NE. We found that Chinmo is expressed in a gradient in the OPC NE that mirrors the expression of Imp, with high levels present at 48 h ALH and declining over time, resulting in the absence of expression at 96 h ALH (Fig. 3F). We observed that Mamo is also temporally expressed in the OPC NE, with its expression initiating 12 hours later than Chinmo, at ∼60 h ALH, and remaining at high levels until 84 h ALH (Fig. 3G).

**Figure 3:**
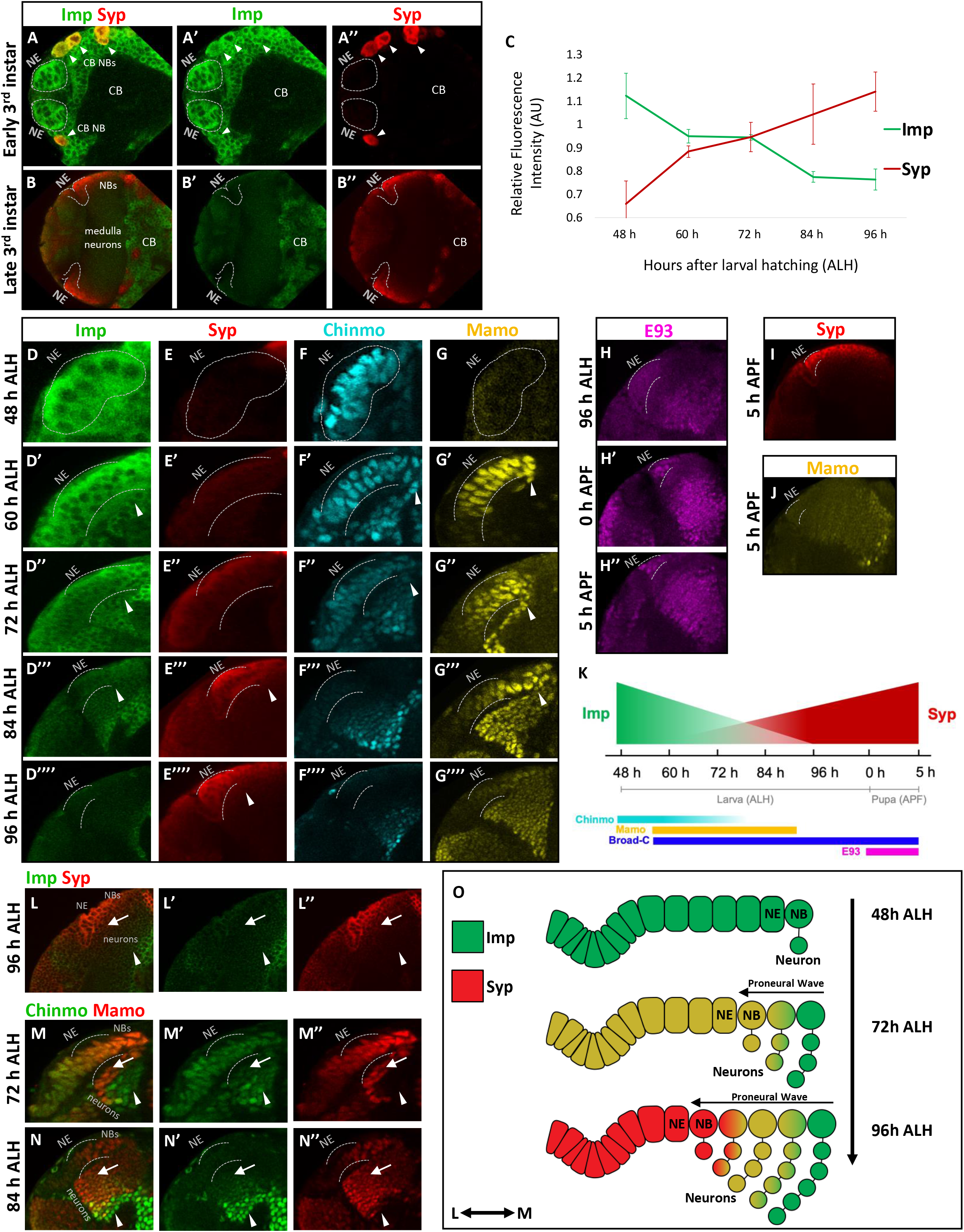
Opposing gradients of Imp and Syp temporally pattern the OPC NE. **A:** Larval brain at the beginning of the third instar. Central brain (CB) is located medial to the optic lobe. Grey dashed lines outline the two arms of the OPC NE. Imp (green) is expressed in the OPC NE, whereas Syp (red) expression is absent. Arrowheads denote CB NBs that co-express high levels of Imp and Syp at this stage. Dorsal is up, medial is right. **B:** Larval brain at the end of the third instar. Central brain (CB) is located medial to the optic lobe. Grey dashed lines outline the two arms of the OPC NE. Imp (green) is expressed at very low levels in the OPC NE, whereas Syp (red) expression levels are high. Dorsal is up, medial is right. **C:** Quantification of Imp (green) and Syp (red) intensities in the OPC NE during third instar larval development. Plot illustrates the mean values of the relative fluorescence intensity for Imp and Syp at each time point (n = 10). **D-G:** Imp (green), Syp (red), Chinmo (cyan) and Mamo (yellow) expression in the OPC at 48, 60, 72, 84 and 96 h ALH. Cross-section views of the OPC with the NE on the left and NBs on the right. Grey dashed lines outline the OPC NE. White arrowheads indicate nascent neurons expressing the respective factors. In the OPC NE, Imp and Chinmo expression decreases, while Syp expression increases, in a gradient manner between 48 h and 96 h ALH. Mamo is strongly expressed in the OPC NE at 60, 72 and 84 h ALH. In all images, medial is right. **H:** E93 (magenta) expression at 96 h ALH (**H**), 0 h (**H’**) and 5 h APF (**H”**) in the OPC NE. Grey dashed lines outline the NE. In the NE, E93 expression begins at 0 h APF and continues to be expressed at 5 h APF. E93 is also broadly expressed in medulla neurons at all three timepoints. **I-J:** Syp (red) and Mamo (yellow) expression at 5 h APF in the OPC NE. Grey dashed lines outline the NE. Syp is expressed at high levels in the NE, whereas Mamo expression is absent. **K:** Summary schematic of the temporal expression of Imp (green), Syp (red), Chinmo (cyan), Mamo (yellow), E93 (magenta) and Broad-C (blue) in the OPC NE over the course of neurogenesis. **L:** 96 h ALH larval brain. Cross-section view of the OPC with the NE on the left and NBs on the right. Imp (green) and Syp (red) expression in the OPC. White arrowhead represents medially located Imp-expressing neurons. White arrow represents laterally located Syp-expressing neurons. **M-N:** Chinmo (green) and Mamo (red) expression in the OPC of 72 h ALH (**M**) and 84 h ALH (**N**) larval brains. Cross-section views of the OPC with the NE on the left and NBs on the right. Grey dashed lines outline the NE. White arrowheads denote medially located Chinmo-expressing neurons. White arrows indicate laterally located Mamo-expressing neurons. **O:** Cartoon schematic depicting how opposing temporal gradients of Imp (green) and Syp (red) assign unique identities to NBs and neurons over the course of neurogenesis. Early-born neurons, generated in the high Imp expression window, are located medially (anterior in the adult), whereas late-born neurons, generated in the high Syp expression window, are located laterally (posterior in the adult).

Our candidate-based approach identified two additional TFs that are temporally regulated in the OPC NE: Br and E93. Br expression initiates in the OPC NE at ∼60 h ALH and remains high through to the end of neurogenesis at 15 h APF (Fig. S3A). E93 is expressed in a late temporal window, initiating at 0 h APF and continuing until the end of neurogenesis (Fig. 3H). During this pupal window, Syp levels remain very high in the NE, whereas the expression of the early TF Mamo is absent (Fig. 3I-J). Taken together, the expression of these TFs in the OPC NE defines four distinct windows of expression: a Chinmo only window from 48-60 h ALH, a Br+Mamo window from 60-84 h ALH, a Br only window from 84 h ALH to 0 h APF and a Broad+E93 window from 0-15 h APF (Fig. 3K). The expression of Chinmo in a declining gradient from 60-84 h ALH may act to further sub-divide the Br+Mamo window. To differentiate these temporally expressed TFs from the tTFs that pattern the NBs as they age, we will here out refer to Chinmo, Mamo, Br and E93 as NE tTFs and the previously described NB factors as NB tTFs.

The NE temporal patterning genes are expressed in medulla NBs and neurons as well (Fig. 3L-N). In NBs, Imp, Syp, and the NE tTFs are expressed in similar temporal windows as observed in the NE. The one exception is Mamo, which is expressed in the OPC NE from 60 h ALH to 84 h ALH, but in a longer window in NBs, from 60 h ALH to 96 h ALH (Fig. 3G). At the neuronal level, Imp and Syp levels correlate to their gradient expression in the NE and NBs, forming complementary gradients of expression in the medulla cortex at 96 h ALH (Fig. 3L).

Chinmo is also expressed in a gradient manner in medulla neurons (Fig. 3M and N). Its expression is highest in the earliest-born, most medial neurons, and decreases in later-born neurons. Mamo is similarly expressed in medial neurons but does not overlap with Chinmo in the earliest-born cells (Fig. 3M and N). Unlike the other NE tTFs, E93 expression in neurons does not mirror its expression in the NE. Rather, it is globally expressed in medulla neurons starting at 96 h ALH and extending through early pupal development (Fig. 3H). This neuronal E93 expression may be regulated by global ecdysone signaling, which has been shown to promote E93 expression during morphogenesis in other tissues (Richards 1976; Baehrecke and Thummel 1995; Lam et al. 2022).

We next determined whether Syp regulates the expression of the NE tTFs. We generated loss-of-function (LOF) *syp* clones and found that Mamo and Chinmo are ectopically expressed in the OPC NE at 96 h ALH, with ectopic expression continuing until the end of larval development (Fig. S3B and C). *syp* is thus required for the repression of Chinmo and Mamo expression in the OPC NE, with its LOF leading to an extension of the Mamo and Chinmo windows from 60-84 h ALH in wild-type brains to 60 h ALH - 0 h APF in mutants. We next analyzed the expression of the late factor E93 in *syp* LOF clones at 5 h APF, a time point when E93 expression levels are high in the wild-type brain (Fig. S3D). We observed that E93 expression is lost in these clones, indicating that *syp* is additionally required to promote E93 expression in the OPC NE. Taken together, these LOF data demonstrate that Syp acts in late temporal windows to repress the expression of the early NE tTFs, Mamo and Chinmo, and promote the expression of the late NE tTF, E93 (Fig. S3E).

The temporal expression of Imp, Syp, and their downstream tTFs in the OPC makes these factors excellent candidates to diversify neuronal fates in the medulla. As the proneural wave sweeps across the NE, newly formed NBs may be assigned unique identities based on the levels of Imp and Syp in the NE cells from which they are derived (Fig 3O). We next asked whether Imp, Syp, and the NE tTFs are required for the specification of Vsx1-Hth neurons.

### Temporal patterning of the OPC NE diversifies Vsx1-Hth neuronal fates

We first determined whether the NE tTFs are expressed in the neurons generated at the intersection of the Vsx1 and Hth axes. At 18 h APF, when neurogenesis is complete, but neurons have not yet redistributed along the D-V axis, we observed that *vsx1⋂hth-Gal4>GFP* cells form a band that extends from medial to lateral in the central region of the medulla cortex (Fig. 4A).

**Figure 4:**
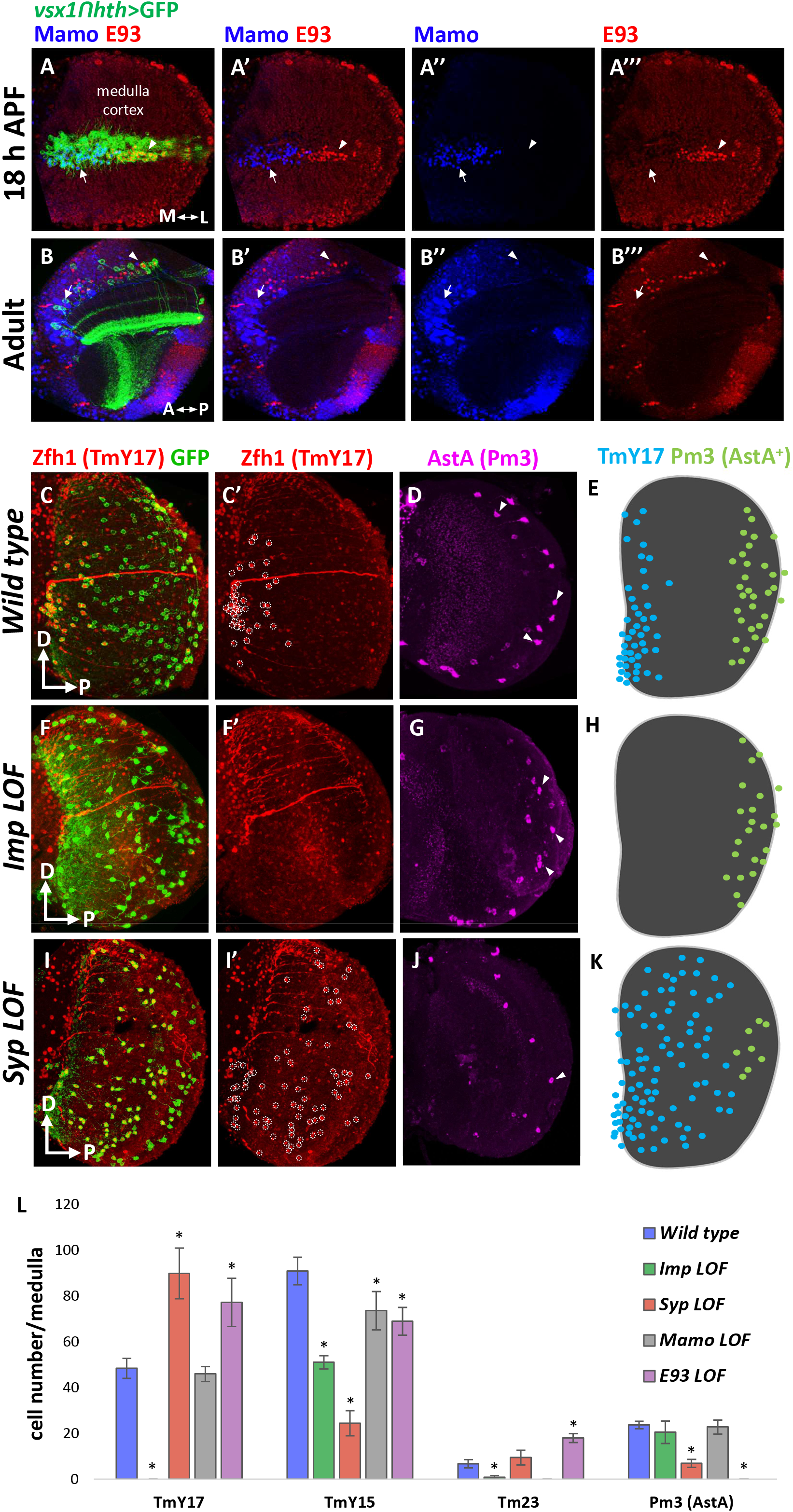
The NE temporal patterning genes regulate Vsx1ꓵHth neuronal fates. **A:** Max projection of *vsx1ꓵhth-Gal4>GFP* (GFP, green) neurons in the medulla cortex at 18 h APF. Within the Vsx1-Hth neuronal band, Mamo (blue) is expressed medially (white arrow), in earlier-born neurons, and E93 (red) is expressed laterally, in later-born neurons (white arrowhead). Medial is left. **B:** Single representative confocal slice of wild type *vsx1ꓵhth-Gal4>GFP* (GFP, green) neurons in the adult medulla. Mamo (blue) is expressed in anteriorly located neurons (white arrow) and E93 (red) is expressed in posteriorly located neurons (white arrowhead). Anterior is left. **C-D:** Max projections of wild-type *vsx1ꓵhth-Gal4>GFP* (GFP, green) neurons and cell-type specific markers in the wild-type adult medulla cortex. (**C**) High levels of Zfh1 expression (red) within *vsx1ꓵhth>GFP* neurons label TmY17 neurons. TmY17 neurons are labeled by high Zfh1 expression (red) and occupy the anterior region of medulla cortex (dashed circles, **C’**). (**D**) AstA (magenta) labels the cell bodies of Pm3a neurons (a subset are indicated with white arrowheads). AstA^+^ Pm3 neurons are located posteriorly. In all images, dorsal is up and posterior is right. **E:** Schematic depicting cell body distributions of TmY17 and Pm3a neurons in the wildtype medulla (n= 12). **F-G:** Max projections of *vsx1ꓵhth>GFP* neurons (GFP, green) and cell-type specific markers in the adult medulla cortex of an *imp* LOF mutant. The *vsx1ꓵhth-Gal4* line was used to drive *UAS*-*imp-*RNAi and *UAS*-*GFP*. (**F**) Zfh1^+^ TmY17 neurons are lost in an *imp* LOF background. (**G**) AstA^+^ Pm3a neurons are still generated in an *imp* LOF background (magenta; a subset are indicated with white arrowheads). **H:** Schematic depicting cell body distributions of TmY17 and Pm3a neurons in an *imp LOF* medulla (n= 9). In *imp* mutants, early-born TmY17 neurons are lost and late-born Pm3a neurons are unaffected. **I-J:** Max projections of *vsx1ꓵhth>GFP* neurons (GFP, green) and cell-type specific markers in the adult medulla cortex of a *syp* LOF mutant. The *vsx1ꓵhth-Gal4* line was used to drive *UAS*-*syp-*RNAi and *UAS*-*GFP*. (**I**) Expansion of Zfh1 expression (red) in *vsx1ꓵhth>GFP* neurons indicates that ectopic TmY17 neurons are generated. Zfh1^+^ TmY17 cell bodies in the *syp* LOF mutant occupy more posterior regions of the medulla cortex (white dashed circles, **I’**). (**J**) AstA^+^ Pm3a neurons (magenta; one is indicated with a white arrowhead) are significantly reduced in number in the *syp* LOF background. **K:** Schematic depicting cell body distributions of TmY17 and Pm3a neurons in a *syp* LOF medulla (n= 17). In *syp* mutants, early-born TmY17 neurons are expanded and late-born Pm3 neurons are reduced. **L:** Graph depicting the mean number of TmY17, TmY15, Tm23 and Pm3a (AstA^+^) neurons in the medulla in wild-type and LOF backgrounds. Asterisks illustrate significant changes in neuron number in LOF versus wildtype (T-test, P<0.05, n= 9-17). Tm23 neurons could not be counted in the *mamo* LOF background because high Mamo expression is the only marker available for Tm23 identification.

These cells represent the progeny of Vsx1-Hth NBs over time, with the earliest-born neurons located medially and the latest-born neurons positioned laterally. We observed that the early NE tTF, Mamo, is expressed in the medial cells of the band, whereas the late NE tTF, E93, is expressed in the lateral portion. The Mamo and E93 expression domains are mutually exclusive, with the expression of the proteins meeting at a boundary two-thirds along the medial-lateral axis. In the adult, Mamo and E93 continue to be differentially expressed in *vsx1⋂hth-Gal4>GFP* neurons along the A-P axis (Fig. 4B). Mamo is expressed in neurons located anteriorly, whereas E93 is expressed in posteriorly localized neurons. Taken together, the above data demonstrate that the temporal patterning of the NE by Mamo and E93 extends to Vsx1-Hth neurons as well.

We next asked whether Imp, Syp, and the NE tTFs are required for the specification of Vsx1-Hth neuronal fates. We used *vsx1⋂hth-Gal4>UAS-RNAi* to knock-down the expression of the temporal genes and analyzed adult brains with cell-type specific markers (Fig. 4C-L). Knock-down of *syp* expression resulted in a dramatic redistribution of Vsx1-Hth neurons from late to early fates in the adult brain; the late neuronal type Pm3 is significantly reduced in number (mean wild-type = 23 compared to mean *syp* LOF = 9, T-test, P<0.05, n= 17), whereas the early cell type TmY17 is almost doubled (mean wild-type = 48 compared to mean *syp* LOF = 89, T-test, P<0.05, n= 17) (Fig. 4C-E and I-L). In contrast, *imp* knock-down resulted in the complete loss of the early-born TmY17 neurons but no change in the number of late Pm3s (Fig. 4C-E, F-H and L). *imp* LOF also resulted in a significant reduction in the number of Tm23 neurons (Fig. 4L). We next analyzed E93, which acts downstream of Syp in the late NE, and found that *e93* knock-down led to a complete loss of Pm3 neurons in the medulla (mean in wild-type = 23 compared to mean in *e93* LOF = 0, T-test, P<0.05, n= 6) as well as the expansion of Tm23 neurons (mean in wild-type = 6 compared to mean in *e93* LOF = 18, T-test, P<0.05, n= 6), which are the cell type born immediately before Pm3 (Fig. 4L, S4A-B, S4E-J). We also observed an expansion of TmY17 neurons, which is surprising given that E93 expression is restricted to the pupal stages in the wild-type OPC NE. We postulate that the global expression of E93 in neurons in the late larva may act to regulate this early fate. Surprisingly, we did not observe changes in the distribution of fates when the early NE tTF Mamo was knocked down, except for a slight decrease in TmY15 neurons (Fig. 4L and Fig. S4A-D). Of note, TmY15 neuronal number is decreased in all LOF backgrounds (Fig. 4L), which suggests that this cell type may require the function of both the early and late temporal patterning factors for its specification. Finally, we were unable to analyze the role of Chinmo in fate specification as *vsx1⋂hth-Gal4>chinmo RNAi* resulted in larval lethality.

Taken together, the above data demonstrate that Imp and Syp are required for the specification of early and late Vsx1-Hth fates, respectively. These findings indicate that, in addition to the previously identified spatial and tTF axes, the temporal patterning of the OPC NE by Imp and Syp represents a third patterning axis that acts to further diversity neuronal fates in the medulla.

### Imp and Syp patterning extends to multiple spatial and temporal windows

Does Imp and Syp patterning extend beyond Vsx1 and Hth to include other temporal and spatial windows? In the temporal axis, we found that the NB tTF series is present at multiple developmental timepoints: Markers for early NB (Exd), intermediate NB (Ey) and late NB (Slp2) windows were observed throughout neurogenesis (Fig. 5A-D). Additionally, we found that the NE tTFs Mamo and Broad are expressed in all medulla NBs at 84 h ALH (Fig. 5E). The expression of the NE tTFs throughout the NB temporal series, together with the observation that the NB tTF cascade is present at multiple developmental stages, suggests that Imp and Syp patterning extends beyond the Hth NB window (Fig. 5F). We next asked whether NB tTF cascade progresses independently of NE patterning by analyzing the progression of the NB temporal series in a *syp* LOF background. We found that *syp* RNAi knockdown driven by *MzVUM-Gal4*, which is expressed in all Vsx1 NE cells and NBs (Erclik et al. 2008), does not affect the progression of the temporal series at 96 h ALH, as the expression of the Exd, Ey, Slp2 and Tll NB tTFs is unaltered (Fig. 5G-I). The above data suggest that the two temporal axes work concurrently and independently to generate neural diversity in the medulla. Finally, in the spatial axis, we observed that Imp, Syp, and the NE tTFs are expressed in NE cells throughout the OPC crescent (Fig 5J-K and S5A-E), which suggests that the temporal patterning of the NE extends beyond the Vsx1 region to include all spatial compartments.

**Figure 5:**
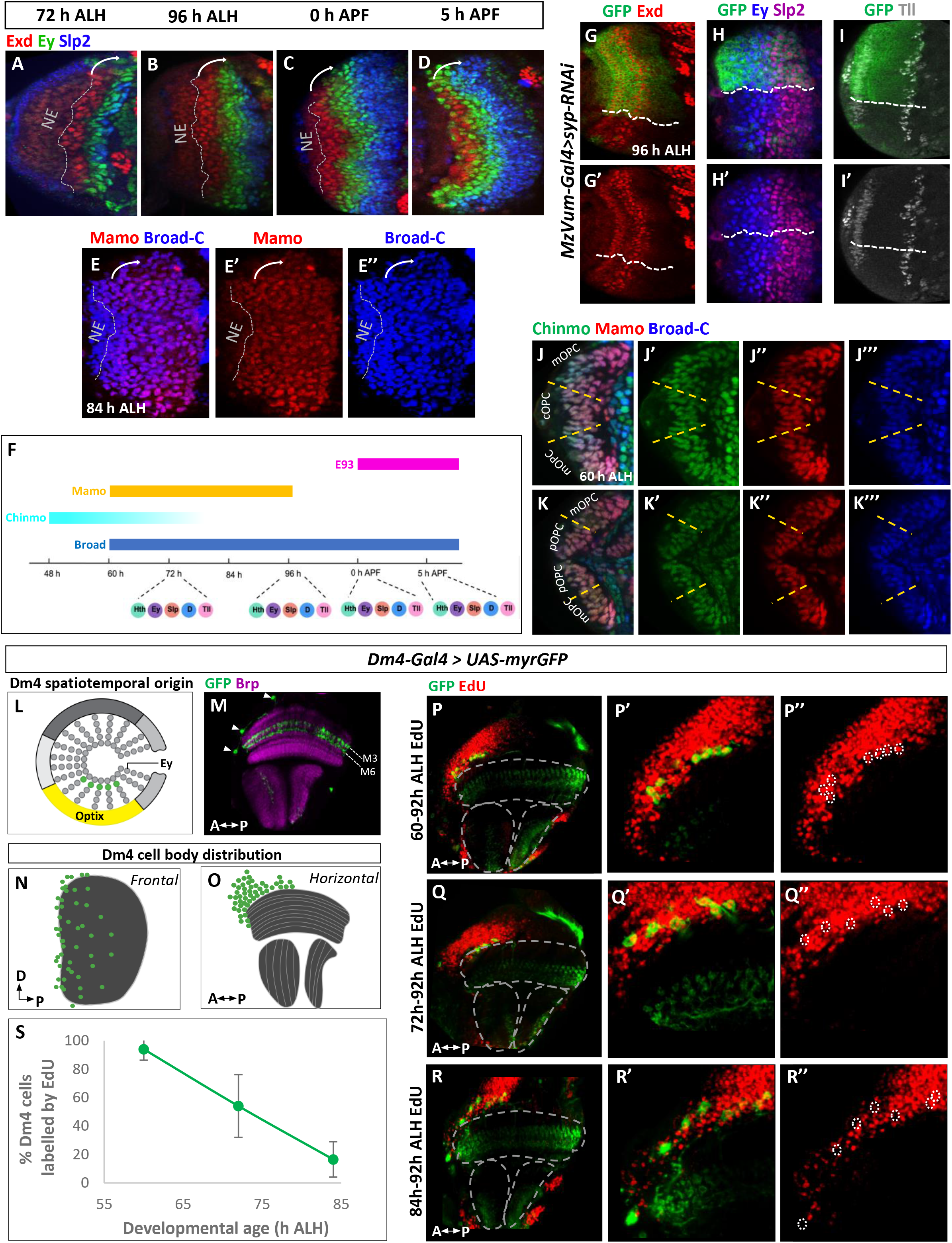
Concurrent temporal patterning generates neural diversity in the medulla. **A-D:** Cortical view of the expression of Exd (red), Ey (green) and Slp2 (blue) in OPC NBs throughout development. (**A**) 72 h ALH, (**B**) 96 h ALH, (**C**) 0 h APF, and (**D**) 5 h APF. Grey dashed line represents the boundary of the OPC NE. White arrow illustrates the progression of the NB temporal sequence. Exd, Ey and Slp2 are expressed in NBs at all larval and pupal timepoints. Dorsal is up, medial is right. **E:** Cortical view of the expression of Mamo (red) and Broad-C (blue) in OPC NBs at 84 h ALH. Grey dashed line represents the boundary of the OPC NE. White arrow illustrates the progression of the NB temporal sequence. Mamo and Broad-C are expressed in all OPC NBs at this development stage. Dorsal is up and medial is right. **F:** Schematic summary of the two temporal patterning mechanisms in the developing optic lobe. **G-I:** Cortical view of the expression of the temporal transcription factors in third instar (96h ALH) *syp* LOF mutants. The *MzVUM-Gal4* line was used to drive *UAS*-*syp-RNAi* and *UAS-GFP* the cOPC. **(G)** Exd (red) labels the first temporal window. (**H**) Ey (blue) and Slp2 (red) label the second and third temporal windows, respectively. **(I)** Tll (grey) labels the last temporal window. Exd, Ey, Slp2, and Tll labeling is unaffected by *syp* knockdown (GFP). Dorsal is up and medial is right. **J:** Expression of Chinmo (green), Mamo (red) and Broad-C (blue) in the cOPC and mOPC of the early third instar (60 h ALH) optic lobe. Chinmo, Mamo and Broad-C expression are continuous in the Vsx1 and Optix spatial domains. Dorsal is up and medial is right. **K:** Expression of Chinmo (green), Mamo (red) and Broad-C (blue) in the pOPC and mOPC of the early third instar (60 h ALH) optic lobe. Chinmo, Mamo and Broad-C expression are continuous in the Rx and Optix spatial domains. Dorsal is up and medial is right. **L:** Schematic illustrating the spatial and temporal origin of Dm4 neurons in the larval OPC. Dm4 neurons are born at the intersection of the ventral Optix spatial domain (yellow) and Eyeless temporal domain (green circles). **M:** Dm4 neurons (GFP; green) in the adult medulla neuropil (magenta) driven by *GMR24F10-Gal4*. Dm4 cell bodies are located in the anterior cortex (arrows in **M**), and innervate layers 3 and 6 of the medulla (dashed lines). **N-O:** Schematic displaying Dm4 cell body positions in both frontal (**N**) and horizontal (**O**) views. Dm4 cell bodies (green) are restricted to the anterior region of the adult medulla (grey). **P-R:** Continuous EdU feeds during larval feeding window to birthdate Dm4 neurons as either early-born or late-born. **(P**) Dm4 neurons are labeled by a 60-92h ALH EdU feed (white dashed circles, **P”**). (**Q**) Dm4 neurons continue to be labeled during a 72-92h ALH EdU feed (dashed circles, **Q”**). (**R**) Very few Dm4 neurons are labeled during an 84-92h ALH feed (dashed circles, **R”**). **S:** Graph showing percent of total Dm4 cells labeled by continuous EdU feeds at different developmental stages. As the onset of continuous EdU feeding is delayed to later time points, fewer Dm4 neurons are labeled. Thus, Dm4 neurons are early-born (n=12 brains per developmental stage, error bars depict standard error of proportions).

To determine whether neurons generated outside of the Vsx1 spatial and Hth temporal axes are also born in restricted developmental windows, we mapped the birthdate of an additional multi-columnar neuron in the medulla, Dm4 (Fig. 5L-M and S5F). We first determined that Dm4 neurons are generated in the ventral Optix region, as memory-trace experiments with the *optix-Gal4* and *hh-Gal4* lines specifically labeled all Dm4 neurons (Fig. S5G-H). Additionally, single-cell RNAseq analysis has placed Dm4 in the intermediate Ey+Homeobrain NB tTF window (Konstantinides et al. 2022). We next plotted the cell body distribution of Dm4 neurons in the adult cortex using a Dm4-specific Gal4-line (*GMR24F10-Gal4*) to drive GFP and found that Dm4 cell bodies are restricted to the anterior region (Fig. 5N-O). This asymmetric cell body distribution of Dm4 suggests that these neurons are born early in development. To birthdate these neurons, we administered EdU to *Dm4-Gal4>GFP* larvae at different time points and found that Dm4 neurons are indeed generated early in development; EdU feeds between 60-92 h ALH and 72-92 h ALH labeled Dm4 neurons (Fig. 5P-Q), whereas feeds between 84-92 h ALH did not (Fig. 5R-S). The early birthdate of Dm4 neurons thus places them in the low Chinmo/Mamo NE tTF window and suggests that Imp and Syp temporal patterning extends beyond Vsx1 and Hth to include the Optix spatial and Ey+Homeobrain temporal windows as well. Taken together, the above data suggest that Imp and Syp patterning of the OPC NE acts as a global mechanism to generate diversity in the medulla.

### Imp and Syp patterning generates unexpected A-P regionalization of the medulla circuit

The restricted cell body positions of Vsx1-Hth neurons along the A-P axis of the medulla cortex led us to investigate whether the arborizations of these neurons innervate all columns of the medulla neuropil. We first analyzed Pm3a innervations using the AstA antibody, which is specifically expressed in Pm3a cell bodies and neurites (Davis et al. 2020). We found that, despite their posteriorly restricted cell bodies (Fig. 2A-B), AstA-labeled neurites cover all medulla columns along the A-P axis (Fig. 6A). Similar results were found with the anteriorly restricted TmY15 neuron (Fig. 2A-B), whose characteristic projections in the distal medulla extend across all columns of the A-P axis in the *vsx1⋂hth-Gal4>GFP* adult brain (Fig. 6B). Our findings for Pm3a and TmY15 are not surprising, since multi-columnar medulla neurons are assumed to innervate all columns in the retinotopic map (Fischbach and Dittrich 1989; Morante and Desplan 2008). One previously reported exception is a sub-type of a Dm8 neuron (DRA-Dm8) that only innervates the columns of the dorsal rim (Courgeon and Desplan 2019; Sancer et al. 2019). Remarkably, however, we found that two Vsx1-Hth neuronal types, TmY17 and TmY12, do not innervate all columns in the medulla, but rather only the columns closest to their cell body location. TmY12 neurons, which are located at the posterior edge of the medulla cortex, only innervate the most posterior medulla columns (Fig. 6C-D, I and S6A-B). Similarly, TmY17 neurons, located in the anterior region of the medulla, send arborizations exclusively to columns in the anterior-ventral region of the medulla neuropil (Fig. 6E-F, J and S6C-D). The striking regionalization of TmY17 and TmY12 innervations indicates that the medulla neuropil contains unanticipated specializations along its A-P axis. It also suggests that Imp and Syp patterning acts to not only generate neuronal diversity in the medulla, but also pattern the retinotopic circuit (Fig. 6L-M).

**Figure 6:**
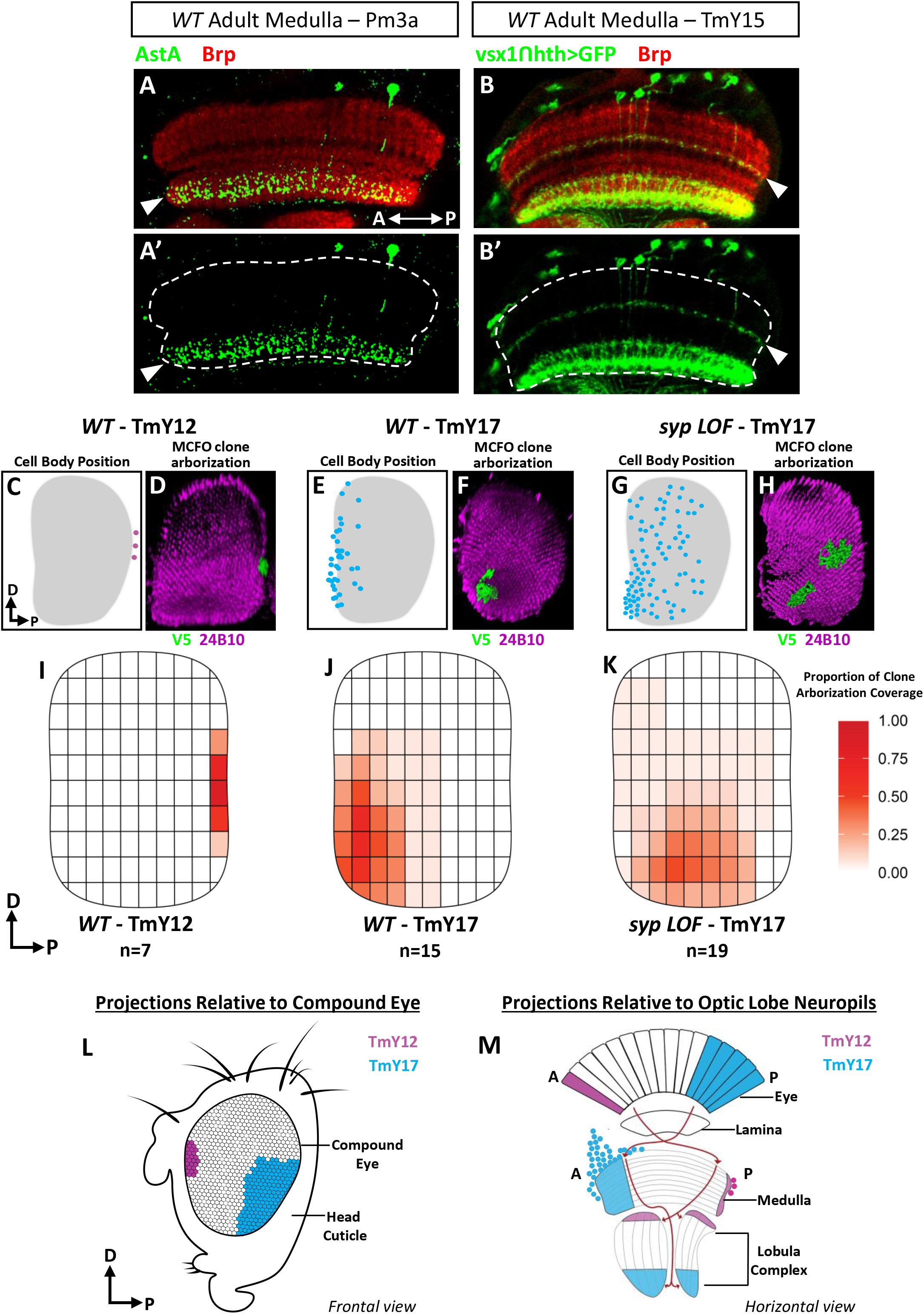
TmY12 and TmY17 neurons innervate distinct columns of the medulla along the A-P axis. **A**: Pm3a projections labeled by AstA (green) innervate the extent of the medulla neuropil (Brp, red) along the A-P axis. White arrowheads indicate Pm3a innervations in the proximal neuropil. Dashed lines (**A’**) outline the medulla neuropil. Posterior is right. **B**: TmY15 projections innervate the extent of the medulla neuropil (Brp, red) along the A-P axis. White arrowhead indicates the distal innervations characteristic of TmY15 labeled by *vsx1ꓵhth-Gal4>GFP* (GFP, green). Dashed lines outline the medulla neuropil (**B’**). Posterior is right. **C, E, G**: Schematics depicting the cell body positions of *vsx1ꓵhth-Gal4* neurons along the D-V and A-P axis of the adult medulla. TmY12 (**C**), wild-type TmY17 (**E**), and TmY17 in a *vsx1ꓵhth-Gal4>syp RNAi* mutant (**G**). Dorsal is up, posterior is right. **D, F, H**: 3D reconstructions of single MCFO-labeled (V5, green) neurons in the adult medulla. Clones of TmY12 (**D**) and TmY17 in a *syp* mutant background (**H**) were generated using *vsx1ꓵhth-Gal4*, while *VT048653ꓵhth-Gal4* was used to generate wild-type TmY17 clones (**F**). 24B10 (Chaoptin, magenta) labels R7/R8 photoreceptor axon terminals and serves as a background stain to visualize medulla columns. Lobula complex arborizations for each neuronal clone were cropped out to better visualize the region of the medulla innervated by the clone along the A-P and D-V axis. Dorsal is up, posterior is right. **I**-**K**: A graphical, heatmap representation of the spatial regions of the adult medulla observed to be innervated by single-cell clones of TmY12 (**I**, n=7), wild-type TmY17 (**J**, n=15) and TmY17 in a *vsx1ꓵhth-Gal4>syp RNAi* mutant (**K**, n=19) neurons. Colour intensity (white-to-red) represents the proportion of total clones observed to innervate a given spatial of the medulla along the A-P and D-V axis (see materials and methods). Dorsal is up, posterior is right. **L**: Schematic illustrating the region of the compound eye subserved by TmY17 (blue) and TmY12 (pink) projections. Frontal view, dorsal is up and anterior is left. **M**: Schematic illustrating the relationship between TmY17 (blue) and TmY12 (pink) projections and their corresponding innervation regions across the optic lobe neuropils. Due to photoreceptor crossover at the first optic chiasm, TmY17 innervations (blue shaded region) process visual information from the posterior region of the overlying eye. TmY12 innervations (pink shaded region) process visual information from the anterior region of the overlying eye. Retinotopic propagation of visual information by TmY17 and TmY12 neurons is also maintained in their projections to the lobula complex.

We next asked whether altering Syp levels leads to a change in the innervation pattern of the locally projecting TmY17 neurons. As noted above, the number of TmY17 neurons is significantly increased (mean in wild-type = 45 to mean in *syp* LOF = 89) in a *syp* LOF background (*vsx1⋂hth-Gal4>syp RNAi*). Consistent with the window of TmY17 production lasting longer in the absence of Syp expression, the cell body positions for these neurons occupy a larger region along the A-P axis, extending into the posterior half of the cortex (Fig. 6G). We generated MCFO clones of *vsx1⋂hth-Gal4>syp RNAi* mutant neurons and mapped the projections of TmY17 neurons (marked by the expression of high Zfh1) onto the retinotopic map. Remarkably, we observed that *syp* LOF TmY17 neuronal projections cover a significantly larger A-P region of the medulla neuropil, extending posteriorly to the same extent as their cell body position (Fig. 6G-H, K and S6E-F). The above data suggest that *syp* is both required for the repression of TmY17 fates and the localization of TmY17 neuron cell bodies (and their projections) to the anterior medulla.

## Discussion

In this study, we identify a third major patterning axis in the OPC that acts together with the previously reported spatial and temporal axes to diversify medulla neuronal fates (Fig. 7). We show that opposing temporal gradients of the Imp and Syp RNA-binding proteins temporally pattern the OPC NE over the three days of medulla neurogenesis. Medulla NBs are thus patterned by concurrent temporal mechanisms: they are assigned a unique temporal identity based on when during development they undergo the NE to NB transition, and they progress through distinct tTF-mediated temporal windows as they age. As a proof of principle, we demonstrate that NBs generated at the intersection of the Vsx1 spatial and Hth temporal axes generate not 1, but 7, distinct neuronal cell types, and that Imp and Syp patterning is required for their specification. We further show that the developmental birthdate of these neurons correlates to their position along the A-P axis in the adult medulla, and that the asymmetric distribution of medulla neurons along this axis leads to unanticipated patterning of the medulla retinotopic circuit. Imp and Syp gradient patterning of the NE thus acts as a powerful mechanism by which to couple neurogenesis with circuit assembly.

**Figure 7:**
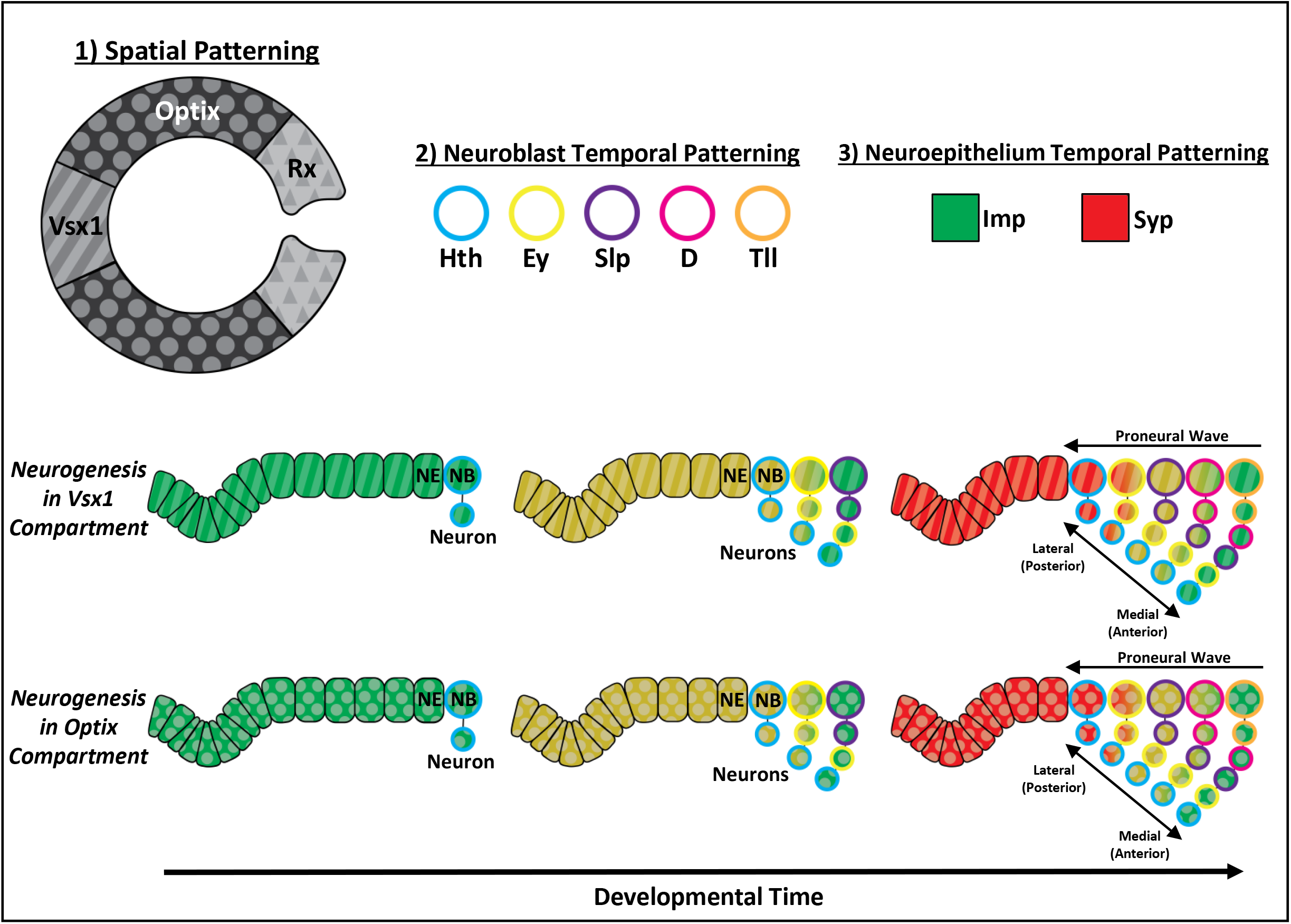
Inputs of three patterning axes in the OPC are integrated to generate neural diversity. Inputs from 3 patterning axes are integrated in neural progenitors of the OPC to generate neural diversity: 1) Expression of spatially expressed factors (Vsx1, Optix and Rx) compartmentalizes the OPC neuroepithelium (NE). 2) Temporally expressed factors (Hth, Ey, Slp, D, Tll) pattern the neuroblasts (NBs) as they age. 3) Temporally expressed RNA-binding proteins (Imp, Syp) pattern the OPC neuroepithelium as it ages.

The identification of a third patterning axis significantly increases the potential amount of neuronal diversity that OPC NBs can generate in the medulla. The combination of 8 spatial compartments, 12 NB temporal windows, and the Notch-based binary diversification of fates can generate up to 192 distinct cell types (Li et al. 2013; Erclik et al. 2017; Konstantinides et al. 2022; Zhu et al. 2022). Based on our findings in the Vsx1-Hth NB window, patterning of the NE by Imp, Syp, and their downstream TFs may increase diversity by another 7-fold. Thus, up to 1344 distinct cell types could theoretically be generated by combinatorial inputs from the three patterning axes and Notch signaling. The programmed cell death of Notch^ON^ or Notch^OFF^ GMC outputs, as observed for neurons in the tOPC region of the medulla and the ventral nerve cord, will likely significantly reduce this number (Lundell et al. 2003; Karcavich and Doe 2005; Bertet et al. 2014). Furthermore, the previous finding that medulla uni-columnar neurons ignore spatial inputs will also lower the number of potential neuronal types generated (Erclik et al. 2017).

However, the identification of a third patterning axis suggests that the previous approximation of 120 neuronal cell types in the adult medulla is an underestimate (Özel et al. 2020). Future studies should extend the analysis of Vsx1-Hth NBs to additional spatio-temporal windows to determine the extent of diversity generated by Imp and Syp patterning. Future research should also determine whether the uni-columnar medulla neurons that ignore spatial patterning inputs are refractory to Imp and Syp patterning as well.

The temporal patterning of the NE, combined with the moving proneural wave that sweeps across the NE (Egger et al. 2007; Yasugi et al. 2008), results in the asymmetric distribution of cell-types along the medial-lateral axis of the larval medulla cortex; neurons generated early in development are located medially, whereas later-born neurons are found laterally. Our finding that Vsx1-Hth neuronal cell types are asymmetrically distributed in the adult cortex suggests that they maintain their medial-lateral positions in the adult. Hence, unlike the extensive neuronal migration previously described across the D-V axis (Erclik et al. 2017), there is only limited migration along the medial (anterior) - lateral (posterior) axis during pupal development.

Additionally, our EdU and memory-trace experiments further demonstrate significantly less movement of cell bodies in the A-P axis. One consequence of limited cell body movement in the A-P axis is that the position of medulla cell types along this axis in the adult can act as a proxy for their developmental birthdate. Indeed, we find that Dm4 neurons are both restricted to the anterior medulla cortex and born in an early developmental window. A second consequence of the asymmetric A-P distribution of neuronal cell types is the unanticipated specialization of the medulla neuropil along this axis. We show that consistent with their localized cell body positions, the innervations of TmY17 neurons are restricted to anterior medulla columns and the innervations of TmY12 neurons are restricted to marginal posterior columns. Future studies should determine whether the A-P patterning of medulla columns plays a role in vision and behavior. Intriguingly, the columns innervated by TmY12 neurons correspond to the ommatidia located at the frontal margin of the compound eye, a region that has been implicated in binocular vision in other insects (Beersma et al. 1977; Seidl and Kaiser 1981; Fischbach and Dittrich 1989; Shinomiya et al. 2019).

Future studies should also employ single-cell RNAseq approaches at multiple timepoints during medulla neurogenesis to identify additional NE temporal patterning genes. The temporal expression of Chinmo, Mamo, Broad, and E93 in the NE can account for four distinct temporal windows. Therefore, it is likely that additional factors play a role in the diversification of fates by this patterning axis. It is anticipated that the genes and mechanisms identified downstream of Imp and Syp in the OPC will also play a role in the central brain and ventral nerve cord, where Imp and Syp temporally pattern both Type I and Type II NB lineages (Liu et al. 2015; Ren et al. 2017; Syed et al. 2017; Liu et al. 2019). Of note, combinatorial temporal patterning has been reported in Type II NBs as well. In the Type II lineage, NBs are temporally patterned by Imp and Syp, and their daughter cells, the intermediate neural progenitors, are patterned by a tTF cascade as they age (Bayraktar and Doe 2013; Ren et al. 2017; Tang et al. 2022). The observation that concurrent temporal patterning mechanisms exist in multiple NB lineages suggests that this patterning strategy may be a common mechanism by which to generate neuronal diversity. Indeed, we anticipate that the concurrent temporal patterning of neural stem cells may play a role in the vertebrate brain as well, where symmetric-to asymmetric-stem cell transitions are common (Cayouette et al. 2006; West et al. 2022).

## Materials and Methods Fly Stocks

Flies were reared at 25ºon standard cornmeal food unless otherwise specified. *OreR* was used as the wild-type strain. The following fly stocks were used in this study: *w*^*1118*^, *vsx1-T2A-Gal4DBD* (this study), *w*^*1118*^*;; hth-T2A-VP16AD* (this study), *w*^*1118*^*;; GMR24F10-Gal4* (*Dm4-Gal4*, BDSC #49090), *w*^*1118*^; *GMR24F10-LexA* (*Dm4-LexA*, BDSC #52696), *MzVum-Gal4* (Erclik et al. 2008), *w*^*1118*^*;; 13xLexAop2-myr::GFP* (BDSC #32212), *w*^*1118*^, *hsFLP;; UAS-FRT>STOP>FRT-myr::smGFP-OLLAS, UAS-FRT>STOP>FRT-myr::smGFP-HA, UAS-FRT>STOP>FRT-myr::smGFP-V5, UAS-FRT>STOP>FRT-myr::smGFP-FLAG* (MCFO-2, BDSC #64086), *w*^*1118*^; *UAS-FRT>STOP>FRT-myr::smGFP-V5, UAS-FRT>STOP>FRT-myr::smGFP-FLAG* (BDSC #62124), *w*^*1118*^, *hsFLPG5::PEST* (BDSC #62118), *UAS-myr::GFP; Bl/CyO; Tm2/Tm6B* (Gift from Larry Zipursky), *hsFlp;; Act5C>CD2>Gal4, UAS-mRFP/ TM3, Sb*^*1*^ (Gift from Dorothea Godt), *w*^*1118*^; *Gal80*^*ts*^; *pxb-Gal4* (Gift from Claude Desplan), *w*^*1118*^; *UAS-Flp; Act-FRT-stop-FRT-nls::βGal* (Gift from Claude Desplan), *UAS-Flp*; *Act-FRT>STOP>FRT-myr::RFP; hh-Gal4* (Gift from Claude Desplan), *UAS-Flp; Optix-gal4; Act-FRT-stop-FRT-nls::βGal* (Gift from Claude Desplan), *w*^*1118*^*;; UAS-Syp-RNAi* (VDRC #33012), *y*^*1*^*sc*v*^*1*^*sev*^*21*^; *UAS-Eip93-RNAi* (BDSC #57868), *y*^*1*^*sc*v*^*1*^*sev*^*21*^; *UAS-Mamo-RNAi* (BDSC #44103), *UAS-Imp-RNAi; UAS-Imp-RNAi/Tm6B* (Gift from Mubarak Hussain Syed), and *w*^*1118*^*;; VT048653-Gal4DBD* (*TmY15-Gal4DBD*, BDSC #73733) (Takemura et al. 2017).

### Immunohistochemistry

Immunohistochemistry analysis on larval and adult brains was performed as previously described (Arain et al. 2021). Immunohistochemistry analysis on pupal brains was performed as described for the adult. The following concentrations of primary antibodies were used to prepare the primary antibody solutions in PBT for a total volume of 100μL: chicken anti-V5 (1:500) (Invitrogen), chicken anti-FLAG (1:200) (Abcam), rabbit anti-HA (1:500) (Invitrogen), chicken anti-GFP (1:1000) (Invitrogen), mouse anti-Brp (nc82, 1:30) (DSHB), rat anti-DE-cadherin (1:20) (DSHB), rat anti-Chinmo (1:500) (Gift from Nicholas Sokol), rat anti DN-cadherin (1:20) (DSHB), guinea pig anti-Vsx1 (1:500) (Erclik et al. 2008), rabbit anti-Hth (1:1000) (Gift from Makoto Sato), mouse anti-Br-core (1:100) (DSHB), mouse anti-Armadillo (1:20) (DSHB), guinea pig anti-Mamo (1:500) (gift from Claude Desplan), mouse anti-Svp (1:200) (DSHB), rabbit anti-E93 (1:300) (Gift from Daniel McKay), rabbit anti-Syp (1:200) (Gift from Claude Desplan), rat anti-Imp (1:200) (Gift from Claude Desplan), rabbit anti-Zfh1 (1:2000) (Gift from Ruth Lehmann), rabbit anti-AstA (1:1000) (Jena Bioscience), guinea pig anti-E93 (1:500) (Gift from Chris Doe), guinea pig anti-Chinmo (1:200) (Gift from Claude Desplan), mouse anti-Chaoptin (24B10, 1:20) (DSHB), goat anti-βgal (1:1000) (MP Biomedicals), rabbit anti-Ey (1:250) (gift from Claude Desplan), guinea pig anti-Slp2 (1:200) (gift from Claude Desplan), mouse anti-Exd (1:200) (DSHB), and guinea pig anti-Tll (1:500) (gift from Claude Desplan).

Secondary antibodies were prepared at 1:500 in PBT for a total volume of 100μL. Secondary antibodies were obtained from Invitrogen and Jackson ImmunoResearch Laboratories.

### Imaging and Image Analysis

Immunostained brains and embryos were imaged on a ZEISS LSM880 confocal microscope using a 25x oil objective. Images were processed using Volocity imaging software or ImageJ.

### Generation of split-Gal4 lines

Vsx1 and Hth split-Gal4 lines were generated via CRISPR-mediated mutagenesis performed by WellGenetics Inc. The split-Gal4 cassettes containing *T2A-VP16AD-RFP* or *T2A-Gal4DBD-RFP* were knocked into the C-terminus of the *vsx1* or *hth* coding region using homology-dependent repair, replacing the stop codon. The gRNAs below were used to target the C-terminus knock-in sites for each gene.

Hth:

> CTCGTAATCCGGCCCCGACC[CGG]

Vsx1:

>GAAGCAGGAGCAGGCGCATC[TGG] / ATCGATGTACATGTTTCATC[TGG]

### Genetic Techniques

#### MCFO clones

MCFO clones of *vsx1* and *hth* co-expressing cells were generated by crossing *w*^*1118*^, *vsx1-T2A-Gal4DBD;;hth-T2A-VP16AD* males to *MCFO-2* females. Adult progeny were heat-shocked for 8 minutes at 37°C and left to recover for 2 days before dissection.

#### TmY17 MCFO Clones

MCFO clones of wild-type TmY17 neurons (Figure 6E, S6C) were generated by crossing virgin females from the previously reported *TmY15-Gal4DBD* (*w*^*1118*^*;; VT048653-Gal4DBD*) (Takemura et al. 2017) to males of *hsFlp;UAS>>smGFP-V5, UAS>>smGFP-FLAG; hth-T2A-VP16AD* virgin females. Adult progeny were heat-shocked for 8 minutes at 37°C and left to recover for 2 days before dissection.

#### Neuroepithelial Syp LOF Clones

To generate *syp* LOF clones, *hsFlp;; Act5C>CD2>Gal4, UAS-mRFP/ TM3, Sb*^*1*^ females were crossed to *w*^*1118*^*;; UAS-Syp-RNAi* males. Larvae were heat-shocked at 37°C for 8 mins. After a two-day recovery, larvae were dissected at the late wandering third instar stage or at 5 h APF.

#### Mutant analysis

To generate LOF mutants, males from each *UAS-RNAi* line (*UAS-syp RNAi, UAS-e93 RNAi, UAS-mamo RNAi*, and *UAS-imp RNAi*) were crossed to *UAS-myr::GFP; Bl/Cyo;Tm2/Tm6B* females. Male progeny were subsequently crossed to *w*^*1118*^, *vsx1-T2A-Gal4DBD;; hth-T2A-VP16AD* females. Adult progeny were dissected and stained post-eclosion. To observe the effect of *syp* LOF on the NB tTFs, *MzVUM-Gal4* females were crossed to *UAS-syp-RNAi* males. The resulting progeny were dissected at the late wandering third instar stage.

### Time course analysis

40-60 OreR flies were raised in bottles at 25°C. To set up an egg lay, parents were transferred to a new bottle with fresh yeast paste. Parent flies were allowed to lay for 3-4 hours before the bottle was cleared. Subsequently, the bottle was incubated at 25°C for 22 hours. The next day, any larvae that hatched prematurely (before 22 hours) were first cleared from the bottle. Fresh yeast paste was added to the bottle and placed back in the 25°C incubator. Newly hatched first-instar larvae (0h ALH) were collected from the bottle at 1-hour intervals.

### Birthdating via EdU

Birthdating experiments within the larval feeding window were performed by administering a nucleotide analog, EdU (5-ethynyl-2’-deoxyuridine) to developing larvae, which is permanently incorporated into newly synthesized DNA. Newly hatched first-instar larvae (0 h ALH) of the genotype, *w*,UAS-myr::GFP/w*^*1118*^,*vsx1-T2A-Gal4DBD;;hth-T2A-VP16AD*, were collected and transferred to EdU-treated food at different developmental stages.

### 48-92h ALH continuous EdU feeds

To make EdU-treated food, 10 mL of standard fly food was melted and poured into a vial. Next, 2 μL of EdU stock solution (50mg/mL) was added for a final concentration of 10μg/mL. After stirring, EdU-treated food was left aside to cool and solidify before use. EdU-treated food can be prepared and stored at 4°C for up to four days. Before transferring larvae to EdU food, refrigerated vials were left at room temperature for up to 1 hour to warm up. After transferring larvae of a specific developmental stage to the EdU food, vials were placed in a 25°C incubator for the remainder of the experiment. Newly hatched adults were immediately transferred from the EdU-treated food to a new, standard food vial.

#### 6-hour EdU pulse feeds

For shorter EdU pulse experiments, 5μL of EdU stock solution (50mg/mL) was added to 10mL of food for a final concentration of 25μg/mL. Red food dye was also added to the food to distinguish larvae that fed on the EdU-treated food from the others. After adding larvae of a specific developmental stage to the food and allowing them to feed for 6 hours at 25°C, the food was scooped out into a dish and only larvae with red dye visible in their midgut were transferred to standard fly food for the remainder of development.

EdU-treated flies were dissected and stained for imaging as described above, with one additional step. After secondary antibody incubation, brain tissues were washed with phosphate-buffered saline (PBS) without Triton overnight, followed by the Click-iT EdU reaction protocol (Thermo Fisher catalog #C10338).

#### EdU cell counts

For each of the 6-hour EdU pulse experiments, the total number of EdU-positive *vsx1ꓵhth-Gal4>GFP* cells was blindly counted first. Next, the EdU-positive cells were sorted into neuronal cell types (TmY17 or TmY15) based on Zfh1 expression to determine the proportion of EdU-positive cells labeled by each cell type at different time points.

### Birthdating via *pxb-Gal4* lineage trace

Neurons born after the larval feeding window (post 92 h ALH) were birth-dated via memory trace experiments using a central OPC driver, *pxb-*Gal4. The presence of a temperature-sensitive Gal80 (Gal80^ts^) in the background facilitated temporal control on *pxb-Gal4* activation at different developmental stages.

Larvae were collected and raised at the permissive temperature (18°C) as described above. Since the average life cycle of *Drosophila* nearly doubles at 18°C (Al-Saffar et al. 1995), developmental age was accounted for at the permissive temperature (i.e. 92h ALH at 25°C corresponds to 184 hours at 18°C). Once at the appropriate age, larvae were transferred to the restrictive temperature of 29°C to inactivate Gal80^ts^ and activate *pxb-Gal4*. Consequently, *pxb-gal4* expression facilitated the removal of a stop cassette by Flp recombinase, leading to constitutive expression of nuclear βgal only in cells generated from the central OPC NE at the time of memory induction.

#### βgal cell counts

Since the *pxb-gal4* lineage labels all cells originating from the central OPC NE, unique marker combinations were used to distinguish Vsx1-Hth neurons among other cell types (TmY17: Zfh1 and Broad, TmY15: Broad and Hth, Tm23: Hth and Mamo, Pm3a: AstA). Total number of cells for each cell type were counted first, followed by the proportion of cells also labeled by βgal.

### Mapping TmY12 and TmY17 Medulla Innervations along the D-V and A-P Axes

Single-cell MCFO-labeled clones of wild-type TmY12, TmY17, and *syp* LOF TmY17 neurons in the adult optic lobe were obtained using the genetic techniques described above. After confocal imaging, 3D rendering and measurement tools on Volocity imaging software were used to measure the A-P and D-V width of each medulla containing a neuronal clone using Brp or NCad staining as a reference. The medulla image was then divided into 10 equally sized spatial regions along the D-V axis and 10 equally sized spatial regions along the A-P axis, forming a 10×10 coordinate grid. The spatial regions of the medulla innervated by each clone were then mapped to their respective medulla coordinate grid along the D-V and A-P axes. For each cell type, the total number of clones innervating a given spatial coordinate was tallied and presented as a proportion of the total number of clones observed. This data was prepared in the form of a heatmap where the intensity of color (white-to-red) reflects the proportion of total clones observed to innervate each spatial region of the medulla.

### Imp and Syp fluorescence analysis

To quantify Imp and Syp expression levels, all larval optic lobes were prepared in a single immunohistochemistry batch. Throughout the time course, Imp and Syp images were acquired using the same confocal settings for each channel. Armadillo (Arm) was used as a counterstain to label the OPC NE. ImageJ was used to make Imp and Syp fluorescence measurements. In a representative slice, hand-drawn sections of the OPC NE (positive integrated density) or the empty background (negative integrated density) were selected on ImageJ using the ROI tool. For each channel, Arm, Imp or Syp, the negative integrated density was subtracted from the positive, resulting in the final integrated density. Within each image, the final Imp or Syp integrated density values were then divided using the final Arm integrated density value, thus resulting in the relative fluorescence intensity value. The relative fluorescence intensity values were then averaged for each timepoint.

#### Analysis of Published Adult Single-Cell RNA-seq Dataset

A published single-cell sequencing dataset on the adult optic lobe (Özel et al. 2020) (GEO accession: GSE142789) was analyzed to identify *vsx1* and *hth* co-expressing cell types. The pre-clustered Seurat object (Adult.rds, GSE142787) was analyzed using the assigned cluster identities located in the “FinalIdents” field of the metadata. Clusters with significantly upregulated *vsx1* or *hth* expression (Padj<0.05, Wilcoxon rank sum test) were identified using the FindAllMarkers function in Seurat 3.1.5 under the following parameters: logfc.threshold = 0, min.pct = 0.3.

Cells within clusters identified to co-express *vsx1* and *hth* were subsetted from the dataset and re-clustered relative to one another using Seurat 3.1.5. All functions were run under default parameters and using 100 principal components (PCs). We chose to perform downstream analysis on clusters resolved at a resolution of 0.8 as this setting cleanly resolved the novel subclusters that had emerged from the re-clustering described above. The FindAllMarkers function was run on the re-clustered data using default parameters to identify unique combinations of marker genes for each of the cell clusters.

## Acknowledgments

We would like to thank Claude Desplan, Tzumin Lee, Chris Doe, Nicholas Sokol, Daniel McKay, Makoto Sato, Mubarak Hussain Syed, Ruth Lehmann, Dorothea Godt, and Larry Zipursky for antibodies and fly stocks. We would like to thank Laurina Manning from the Doe lab for her ongoing assistance with EdU birth-dating protocols. Stocks obtained from the Bloomington Drosophila Stock Center (NIH P40OD018537) and Vienna Drosophila Resource Center were used in this study. The DE-Cad, NCad, Svp, Brp, Chaoptin, Arm, Br-core, and Exd monoclonal antibodies were obtained from the Developmental Studies Hybridoma Bank, created by the NICHD of the NIH and maintained at The University of Iowa, Department of Biology, Iowa City, IA 52242. Schematics were generated using Adobe Illustrator.

## Funding

This work was supported by an NSERC Discovery Grant (RGPIN2015-06457) awarded to T.E. P.V. is supported by the Vision Science Research Program (University of Toronto: Ophthalmology and Vision Sciences and UHN), the Ontario Graduate Scholarship, and the Queen Elizabeth II/Pfizer Graduate Scholarship in Science and Technology. U.A is supported by the NSERC Alexander Graham Bell Canada Graduate Scholarship (CGSD-518890-2018).

## Figure Captions

**Figure 1 Supplement:**
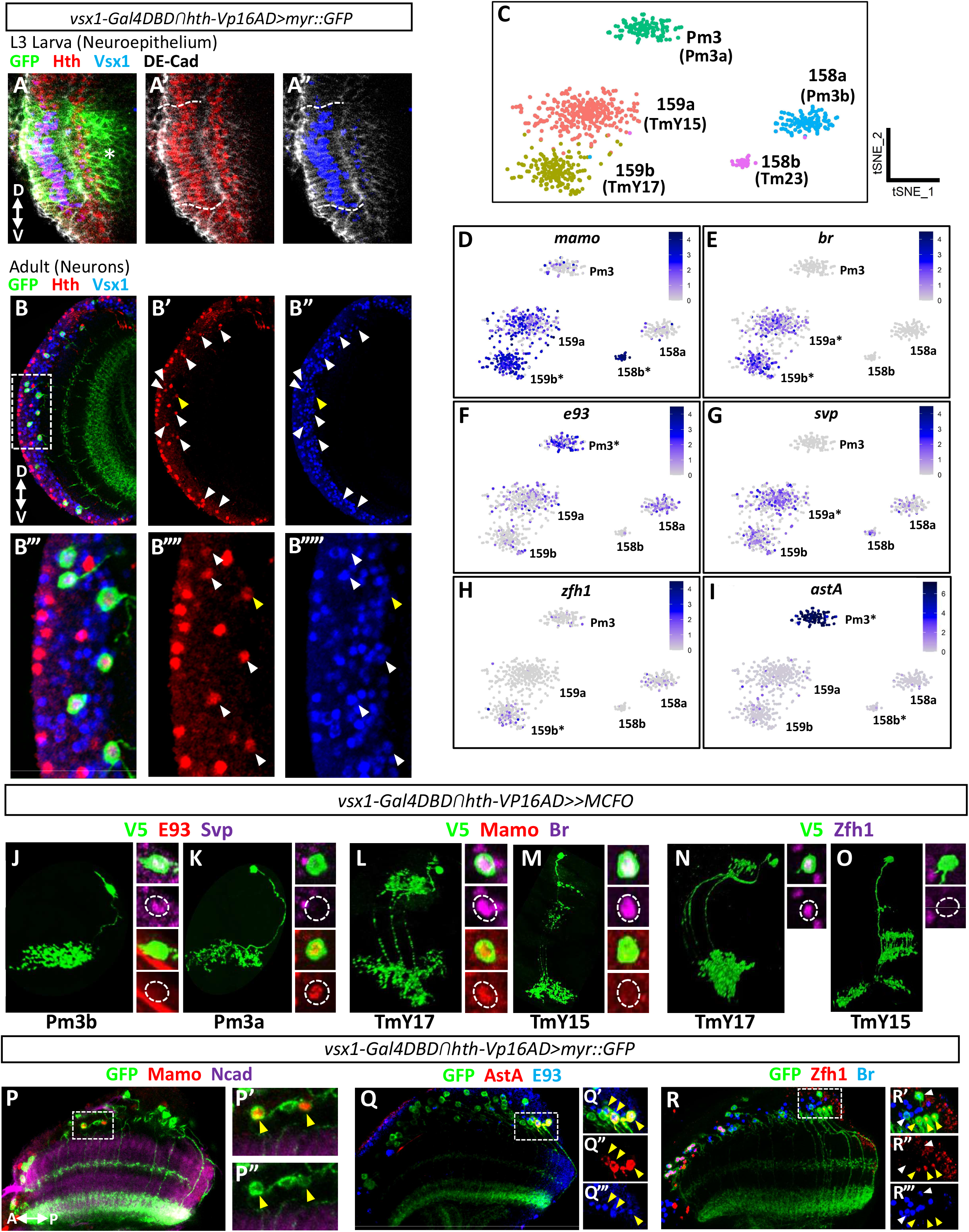
Identification of markers that differentially label vsx1ꓵhth neurons. **A**: **A**: GFP (green) expression in OPC NE (DE-Cad) of a L3 larva as driven by *vsx1ꓵhth-Gal4*. Hth (red) and Vsx1 (blue) are co-expressed in the GFP-labeled NE cells. Dashed lines outline the boundaries of active GFP expression in the OPC NE and the asterisk marks the region of GFP signal resulting from perdurance in neurons. Dorsal is up. **B**: Hth (red) and Vsx1 (blue) are co-expressed in neurons labeled by GFP (green) driven by *vsx1ꓵhth-Gal4* in the adult medulla cortex (arrowheads). The neuron indicated with a yellow arrowhead expresses low levels of Vsx1. The cortex region outlined with dashed lines (**B**) is magnified in the panels below (**B’’’-B’’’’’**). Dorsal is up. **C**: tSNE visualization of *vsx1* and *hth* co-expressing cells subsetted from a whole adult *Drosophila* optic lobe single-cell RNAseq dataset. Neuronal clusters are annotated with their published cluster identities as well as new cell type identities assigned in accordance with marker analysis performed on the *vsx1ꓵhth-Gal4* line. **D-I**: tSNE plots of *vsx1* and *hth* co-expressing cells colored by their non-integrated expression level of *mamo, br, e93, svp, zfh1*, and *astA*. Asterisks indicate the cell clusters with significantly higher expression of the respective gene relative to the other clusters (Wilcoxon Rank Sum, P-adj<0.05). **J**-**K:** E93 (red) and Svp (magenta) expression in *vsx1ꓵhth-Gal4* MCFO clones (V5, green) of Pm3b (**J**) and Pm3a (**K**) neurons. Max projection images of the neuronal clones are on the left and single confocal sections showing marker expression in the cell bodies (dashed lines) are on the right. **L-M**: Mamo (red) and Br (magenta) expression in *vsx1ꓵhth-Gal4* MCFO clones (V5, green) of TmY17 (**L**) and TmY15 (**M**) neurons. Max projection images of the neuronal clones are on the left and single confocal sections showing marker expression in the cell bodies (dashed lines) are on the right. **N-O**: Zfh1 (magenta) expression in *vsx1ꓵhth-Gal4* MCFO clones (V5, green) of TmY17 (**N**) and TmY15 (**O**) neurons. 3D reconstructions of the neuronal clones are on the left and single confocal sections showing marker expression in the cell bodies (dashed lines) are on the right. **P**: Mamo (red) expression in *vsx1ꓵhth-Gal4>GFP* neurons (GFP, green) in the adult medulla (Ncad, magenta). The medulla cortex region outlined with dashed lines (**P**) is magnified in the adjacent panels (**P”-P’’’**). Yellow arrowheads indicate Tm23 cell bodies which possess high Mamo expression and reside in the proximal cortex. Anterior is left. **Q**: AstA (red) and E93 (blue) are co-expressed in a subset of *vsx1ꓵhth-Gal4>GFP* neurons (GFP, green). The cortex region outlined with dashed lines (**Q**) is magnified in the adjacent panels (**Q’-Q’’’**). All Pm3a neurons (E93^+^, yellow arrowheads) express AstA. Anterior is left. **R**: Zfh1 (red) and Br (blue) expression in *vsx1ꓵhth-Gal4>GFP* neurons (GFP, green). The cortex region outlined with dashed lines (**R**) is magnified in the adjacent panels (**R’-R’’’**). All Pm3 neurons (Br ^-^, yellow arrowheads) express low levels of Zfh1. White arrowheads indicate TmY15 neurons (Br^+^). Anterior is left.

**Figure 2 Supplement:**
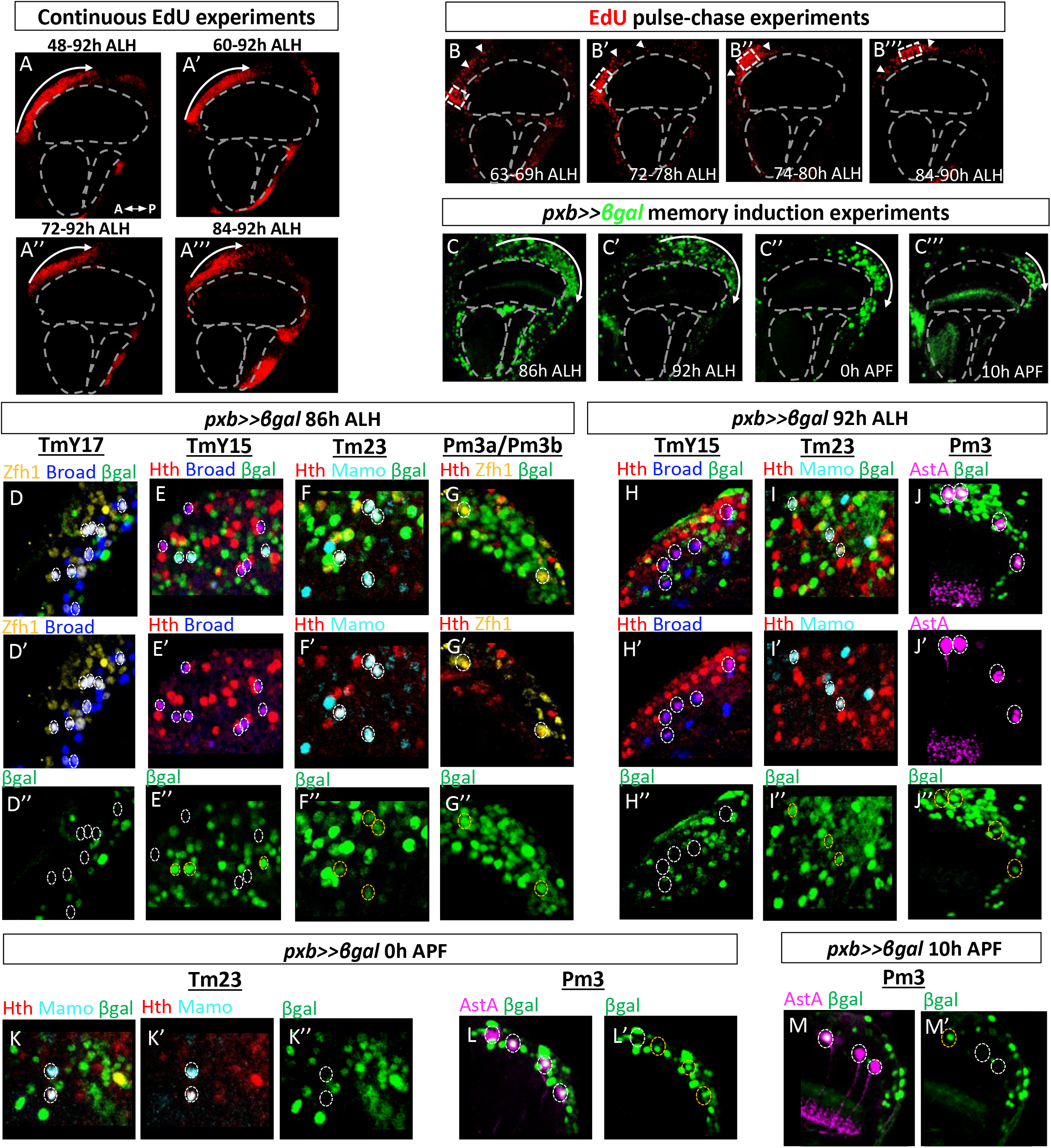
Lineage-based clonal experiments reveal the sequential birth-order of TmY15, Tm23 and Pm3 neurons. **A**: Representative images of continuous EdU feeds starting at distinct time points within the feeding window. (**A**) EdU feed starting at 48 h ALH labels the full extent of early-born cells in the anterior half of the adult medulla cortex (red). (**A’-A’’’**) As the onset of continuous EdU feeding is delayed to later time points, the extent of cells labeled in the anterior cortex is reduced, but the termination point (∼92 h ALH, center of A-P axis) remains unchanged. **B**: Representative images of 6-hour EdU pulse experiments within the feeding window. (**B**) EdU pulse in the early third-instar (63-69 h ALH) forms a band that labels the most anterior cells in the medulla cortex (dashed rectangle). (**B’-B”**) EdU pulse in mid third-instar (72-78 h ALH or 74-80 h ALH) labels cells adjacent to those labeled in (**B**) (dashed rectangle). (**B’’’**) EdU pulse in the late third-instar (84-90 h ALH) labels most posterior cells in the feeding window (dashed rectangle). White arrows label cells that migrated away from the band along the A-P axis at each time point. Notably, cell dispersion in the A-P axis is limited, indicating that the relative A-P position of cells in the medulla cortex indicates the developmental stage during which they were generated. **C**: Memory induction experiments using a central OPC NE (*vsx1* region) driver, *pxb>Gal4*. Memory induction via heat shock was utilized to birthdate cells that could not be labeled by EdU feeds. (**C-C’**) Memory induction at late third-instar (86 or 92 h ALH) labels only posterior cells in the adult medulla cortex. (**C”-C’’’**) Memory induction in pupal stages labels the most posterior cells of the medulla cortex (white arrows). As the onset of memory induction is delayed to later time points, the extent of cells labeled in the cortex is reduced, but the relative location of labeled cells shifts more posteriorly. **D-G**: Heat shock at 86 h ALH does not label early-born TmY17 neurons, but does label TmY15, Tm23, and Pm3 neurons. (**D-D’’’**) Zfh1^+^, Br^+^ TmY17 cells are not labeled by βgal memory (dashed circles). (**E-E”**) Br^+^, Hth^+^ TmY15 neurons are labeled by βgal, but not entirely (white dashed circles label βgal^-^ cells, while yellow dashed circles label βgal^+^ cells). (**F-F”**) Hth^+^, Mamo^+^ Tm23 neurons are labeled by βgal (yellow dashed circles). (**G-G”**) Weakly Zfh1^+^, Hth^+^ Pm3 neurons are also labeled by βgal (yellow dashed circles). **H-J**: Heat shock at 92 h ALH no longer labels TmY15 neurons, but continues to label Tm23 and Pm3 neurons. (**H-H”**) Br^+^, Hth^+^ TmY15 neurons are no longer labeled by βgal memory (white dashed circles), whereas Hth^+^, Mamo^+^ Tm23 neurons (**I-I”**) are labeled (yellow dashed circles). (**J-J”**) Pm3a neurons, marked by AstA are also labeled by a 92 h ALH memory induction (yellow dashed circles). **K-L**: Heat shock at 0 h APF no longer labels Tm23 neurons, but continues to label Pm3 neurons. (**K-K”**) Hth^+^, Mamo^+^ Tm23 neurons are no longer labeled (white dashed circles). (**L-L’**) AstA^+^ Pm3 neurons continue to be labeled by βgal memory at 0h APF (yellow dashed circles). **M**: Heat shock at 10 h APF labels only a few Pm3 neurons (yellow dashed circles).

**Figure 3 Supplement:**
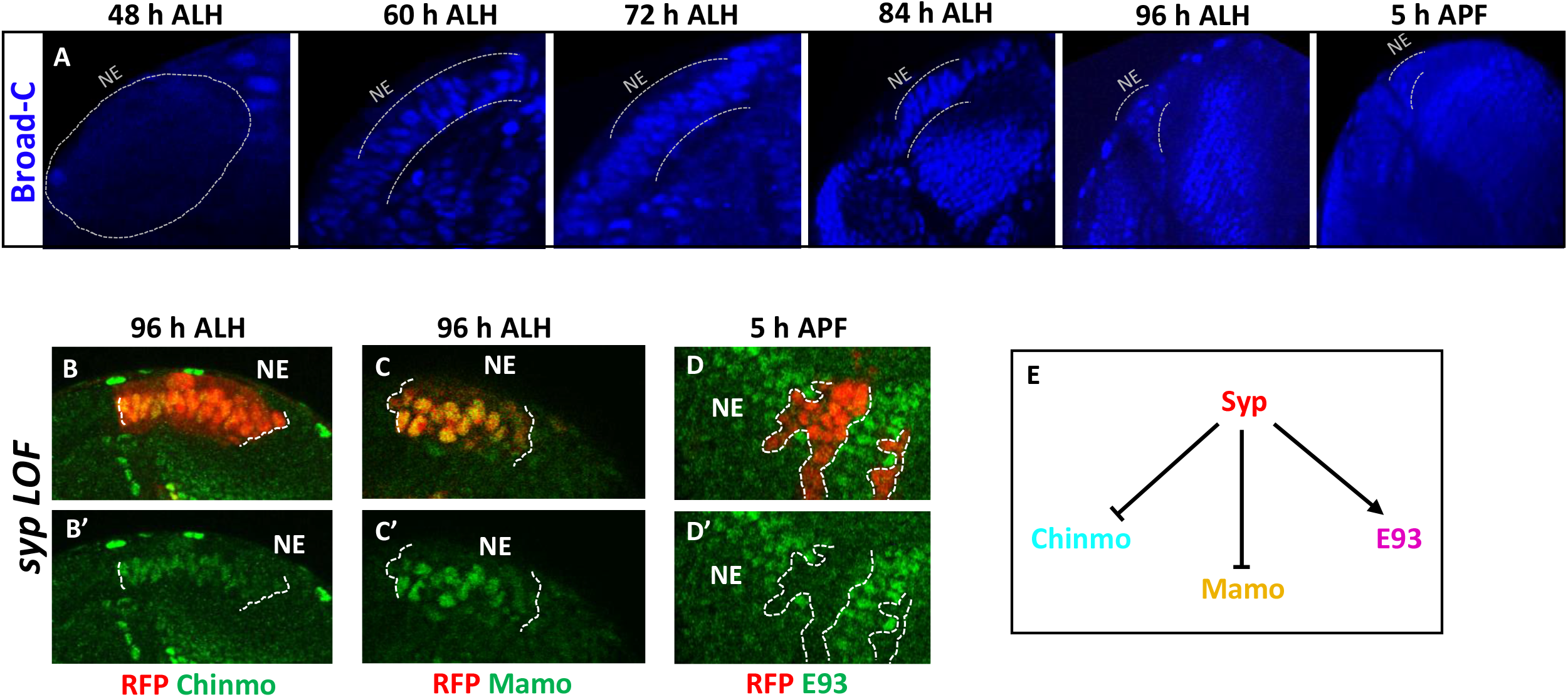
Syp regulates the early factors Mamo and Chinmo, and the late factor E93 in the OPC NE. **A:** Br (blue) expression at 48, 60, 72, 84, 96 h ALH and 5 h APF in the OPC NE. Grey dashed lines outline the NE. **B:** Upregulation of Chinmo expression (green) in *syp* RNAi LOF clones (RFP, red) in the OPC NE at 96 h ALH. White dashed lines outline the boundary of the clone. **C:** Upregulation of Mamo expression (green) in *syp* RNAi LOF clones (RFP, red) in the OPC NE at 96 h ALH. White dashed lines outline the boundary of the clone. **D:** Loss of E93 expression (green) in *syp* RNAi LOF clones (RFP, red) in the OPC NE at 5 h APF. White dashed lines outline the boundaries of the clones. **E:** Regulatory relationships between Syp and the downstream factors Chinmo, Mamo and E93.

**Figure 4 Supplement:**
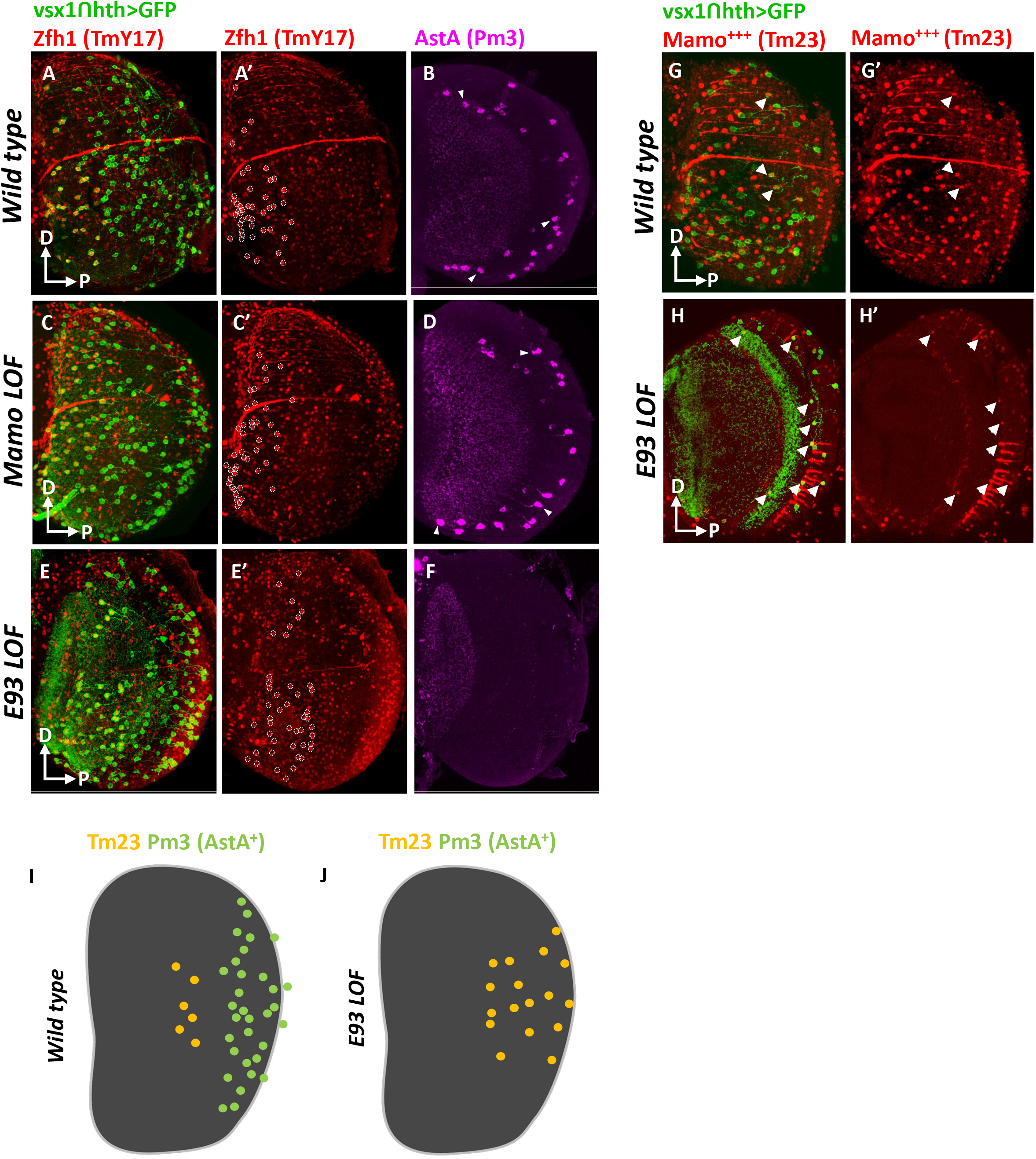
The Mamo and E93 windows contribute to neuronal diversity within the Vsx1ꓵHth spatio-temporal domain. **A-B:** Max projections of wild-type *vsx1ꓵhth-Gal4>GFP* neurons (GFP, green) and cell-type specific markers in the wild-type adult medulla cortex. (**A**) TmY17 neurons are labeled by high Zfh1 expression (red) and occupy the anterior region of medulla cortex (dashed circles, **A’**). (**B**) AstA (magenta) labels the cell bodies of Pm3a neurons (a subset of which are indicated by white arrowheads). Pm3a neurons are located posteriorly. **C-D:** Max projections of *vsx1ꓵhth>GFP* neurons (GFP, green) and cell-type specific markers in the adult medulla cortex of a *mamo* LOF mutant. The *vsx1ꓵhth-Gal4* line was used to drive *UAS*-*mamo-*RNAi and *UAS*-*GFP*. (**C**) Zfh1 expression (red) within *vsx1ꓵhth>GFP* neurons is comparable to wild-type, indicating that TmY17 neurons (dashed circles, **C’**) are unaffected. (**D**) AstA (magenta) labels the cell bodies of Pm3a neurons (a subset of which are indicated with white arrowheads). Pm3a neurons are unaffected in the *mamo* LOF background. **E-F:** Max projections of *vsx1ꓵhth>GFP* neurons (GFP, green) and cell-type specific markers in the adult medulla cortex of a *e93* LOF mutant. The *vsx1ꓵhth-Gal4* line was used to drive *UAS*-*e93-*RNAi and *UAS*-*GFP*. (**E**) Zfh1 expression (red) within v*sx1ꓵhth>GFP* neurons indicates TmY17 neurons (dashed circles, **E’**). (**D**) AstA (magenta) labels the cell bodies of Pm3a neurons. Pm3a neurons are entirely lost in *e93* LOF mutants. **G:** Max projections of *vsx1ꓵhth-Gal4>GFP* (green) neurons and cell-type specific markers in the wild-type adult medulla cortex. High levels of Mamo expression (red) within *vsx1ꓵhth-Gal4>GFP* neurons label Tm23 neurons (white arrows). Tm23 neurons are located in the center of the A-P axis. **H:** Max projections of *vsx1ꓵhth>GFP* neurons (GFP, green) and cell-type specific markers in the adult medulla cortex of a *e93* LOF mutant. The *vsx1ꓵhth-Gal4* line was used to drive *UAS*-*e93-*RNAi and *UAS*-*GFP*. Mamo (red) within *vsx1ꓵhth-Gal4>GFP* neurons indicate Tm23 neurons (white arrows). White arrows represent the distribution of Tm23 neurons in the *e93* LOF mutant optic lobe. Tm23 neurons increase in number and are located more posteriorly in *e93* LOF mutants. **I:** Schematic depicting cell body distributions of Tm23 and Pm3a neurons in the wild-type medulla (n= 6). **J:** Schematic depicting cell body distributions of Tm23 and Pm3a neurons in the *e93* LOF medulla (n= 6). In *e93* mutants, Tm23 neurons increase in number and are shifted posteriorly while Pm3a neuronal fates are lost. In all images: Dorsal is up, posterior is right.

**Figure 5 Supplement:**
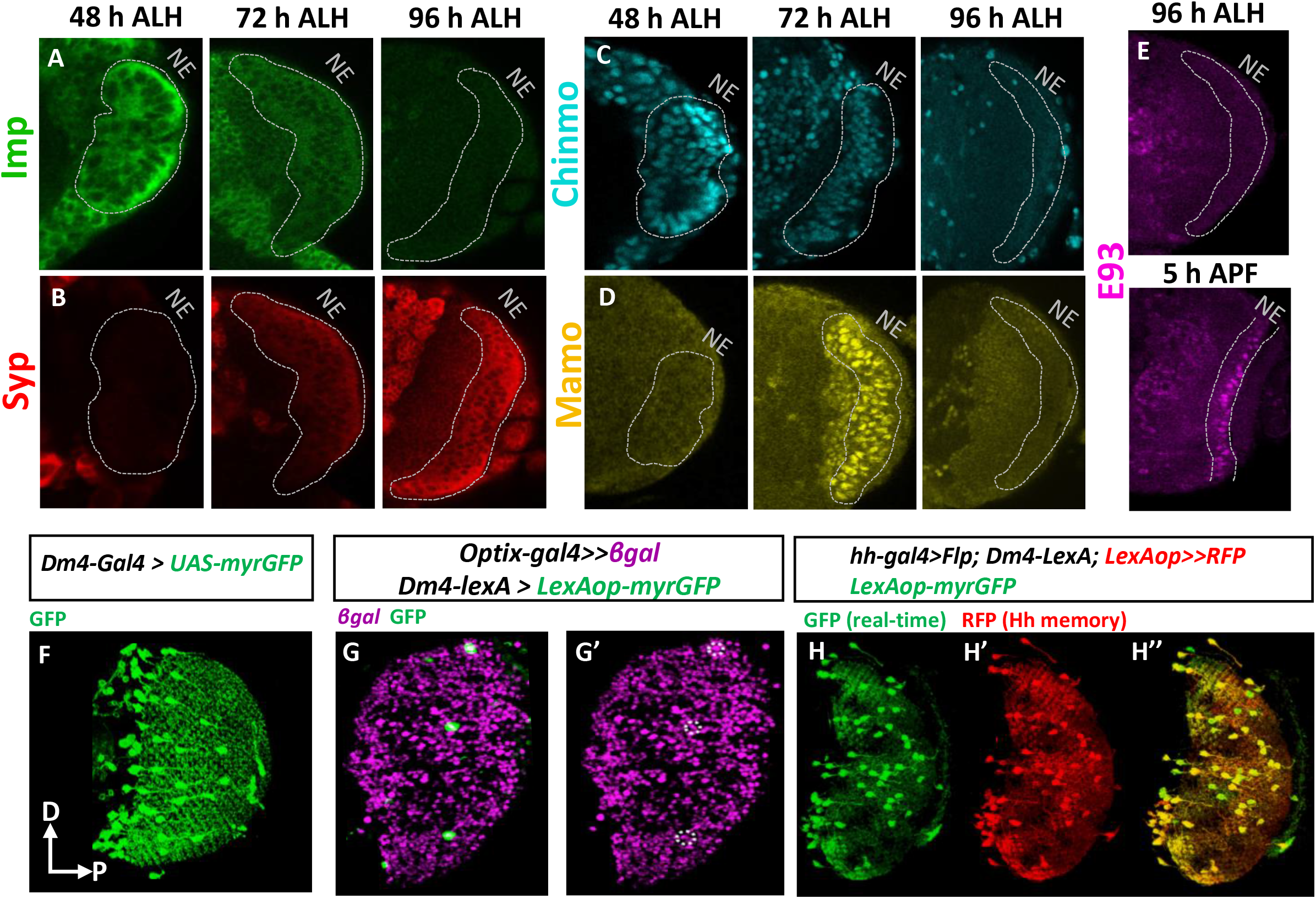
Concurrent temporal patterning mechanisms extend across all spatial compartments of the OPC. **A-D:** Expression of Imp (green), Syp (red) and Chinmo (cyan) and Mamo (yellow) in the cOPC and mOPC throughout development. Expression of NE temporal factors are continuous in all spatial compartments. **E:** Expression of E93 (magenta) in the cOPC and mOPC at 96 h ALH and 5 h APF. At 5 h APF, expression of E93 is continuous in all spatial compartments. **F:** 3D projection of Dm4 neurons labeled by GFP in the frontal orientation. Dorsal is up and posterior is right. Cell bodies of Dm4 neurons (green) are restricted to the anterior medulla cortex, but innervations cover the entire A-P and D-V axes of the medulla. **G:** Optix lineage-trace with nuclear βgal (magenta) labels all mOPC-derived neurons in the adult medulla cortex. Dm4 neurons labeled by GFP (green) are also co-labeled by βgal (dashed circles), indicating that Dm4 neurons derive from the Optix spatial domain. **H:** *hh* lineage trace in combination with the *Dm4-lexA* driver indicates that Dm4 neurons are derived from the ventral OPC. Stop-cassette excision in *LexAop>Stop>RFP* is only mediated in ventral OPC cells as a result of *hh-Gal4>Flp*. Thus, all Dm4s alabeledled by *Dm4-lexA>GFP* (GFP, green), but only ventrally derived Dm4s are labeled by *Dm4-LexA>RFP* (RFP, red). As all Dm4s are labeled by both GFP and RFP, Dm4s are exclusively derived from the ventral mOPC.

**Figure 6 Supplement:**
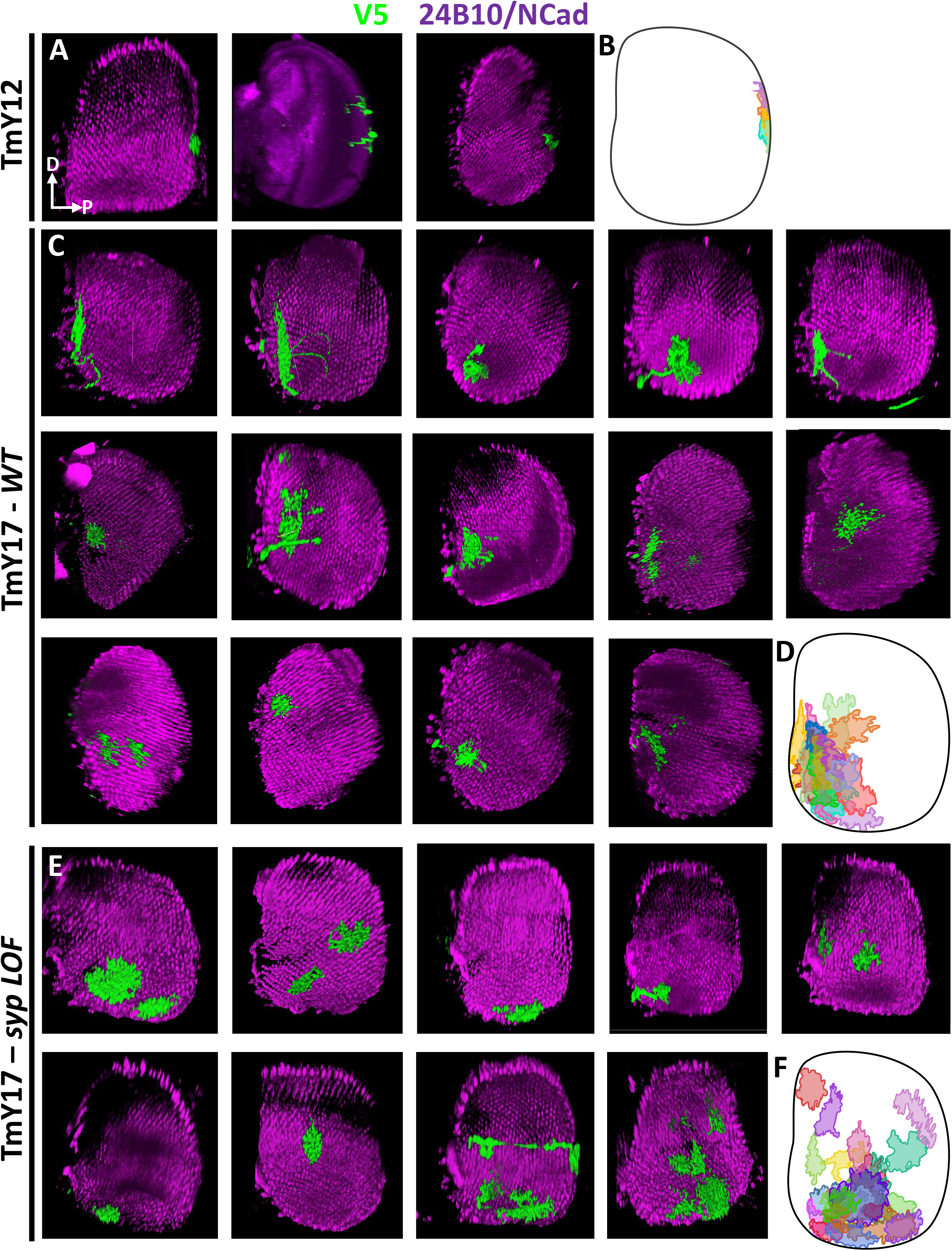
Innervations of single TmY12 and TmY17 clones in the medulla. **A, C, E**: 3D reconstructions of single MCFO-labelled (V5, green) neurons in the adult medulla Ncad or 24B10/Chaoptin, magenta). Clones of TmY12 (**A**) and TmY17 in a *syp* mutant background (**E**) were generated using *vsx1ꓵhth-Gal4*, while *VT048653ꓵhth-Gal4* was used to generate wild-type TmY17 clones (**C**). Lobula complex arborizations for each neuronal clone were cropped out to better visualize the region of the medulla innervated by the clone along the A-P and D-V axis. In all images dorsal is up, posterior is right. **B, D, F**: Schematic traces of the medulla arborizations of single-neuron MCFO clones (a subset of which are traced from the clones displayed in **A, C**, and **E**) overlayed onto a schematic representation of a medulla. TmY12 (**B**), wild-type TmY17 (**D**), and TmY17 in a *vsx1ꓵhth-Gal4>syp RNAi* mutant (**F**). In all images dorsal is up, posterior is right.

## Notes

### Competing Interest Statement

The authors have declared no competing interest.

